# Artificial intelligence as a ploy to delve into the intricate relationship between genetics and mitochondria in MASLD patients

**DOI:** 10.1101/2024.06.03.597155

**Authors:** Miriam Longo, Erika Paolini, Marica Meroni, Michela Ripolone, Laura Napoli, Francesco Gentile, Annalisa Cespiati, Marco Maggioni, Anna Alisi, Luca Miele, Giorgio Soardo, Maurizio Moggio, Anna Ludovica Fracanzani, Paola Dongiovanni

## Abstract

**Background and Aims:** Mitochondrial (mt-) dysfunction is a hallmark of progressive MASLD. MtDNA copy number (mtDNA-CN) and cell-free circulating mtDNA (ccf-mtDNA), which reflect mt-mass and mt-dysfunction, respectively, are gaining attention as non-invasive disease biomarkers. We previously demonstrated that *PNPLA3*/*MBOAT7*/*TM6SF2* deficiency in HepG2 cells increased mt-mass, mtDNA-CN and ccf-mtDNA. This study furtherly explored mt-biogenesis, function and mt-biomarkers in biopsied MASLD patients from a Discovery (n=28) and a Validation (n=824) cohort, stratified by the number of risk variants (NRV=3). We took advantage of artificial intelligence (AI) to develop new risk scores, predicting MASLD evolution by integrating anthropometric and genetic data (Age, BMI, NRV) with mt-biomarkers.

**Methods:** Hepatic mt-morphology and dynamics were assessed by TEM, IHC and gene expression. mtDNA-CN and ccf-mtDNA were measured in PBMCs and serum samples. GPT-4 was employed as AI tool to support the construction of novel risk scores for MASLD progressive forms (MASH, fibrosis and HCC).

**Results:** In the Discovery cohort, NRV=3 patients showed the highest mt-mass and significant mt-morphological changes (i.e. membranes rupture). An elevated PGC-1α, OPA1, DRP1 and PINK1, markers of mt-biogenesis, fusion and fission were found in these patients, supporting an enhanced mt-dynamics. However, PRKN protein levels were reduced, suggesting a premature block of mitophagy. In the Validation cohort, *PGC-1α* mRNA levels and mtDNA-CN were significantly higher in NRV=3 compared to patients with 1,2 or no variants. Circulating mtDNA-CN and ccf-mtDNA were augmented in NRV=3 patients and correlated with genetics and MASLD severity at multivariate analysis, supporting that both may independently modulate mt-dynamics and activity. By exploiting rsGPT-4 we then optimized the combination of non-invasive variables to get prediction models named Mitochondrial, Anthropometric, and Genetic Integration with Computational intelligence (“MAGIC-“) for assessing MASH, fibrosis, and HCC, respectively. The MAGIC-MASH and MAGIC-Fib models showed AUCs of 73% and 76% in detecting MASH and fibrosis >1. Of note, MAGIC-HCC achieved an AUC of 86% (95% CI: 0.823-0.885), with 78.6% sensitivity and 81.5% specificity thus resulting the best score for the desired outcome.

**Conclusions:** mtDNA-CN and ccf-mtDNA may have pathological and prognostic significance in MASLD patients, especially in those genetically-predisposed.

## Introduction

Metabolic-dysfunction associated steatotic liver disease (MASLD) spans from steatotic liver disease (SLD) to metabolic dysfunction-associated steatohepatitis (MASH), fibrosis, cirrhosis, and hepatocellular carcinoma (HCC) (1). Three missense polymorphisms (SNPs) in *PNPLA3*, *MBOAT7* and *TM6SF2* genes, which regulate lipid metabolism, have been recognized as the major MASLD genetic predictors. Moreover, the cumulative weight of these mutations increases the risk of presenting a higher histological damage at the time of diagnosis and to progress up to HCC (2–7).

Mitochondria are dynamic organelles toggling between fusion, fission and mitophagy events in a balanced manner, the latter occurring to remove damaged mitochondria. In response to endogenous/exogenous *stimuli*, mitochondria adapt their morphology, number and activities to maintain cellular homeostasis. During MASLD progression, this adaptability is lost, leading to the formation of unfunctional and misshaped mitochondria (8–10). Although the role of mt-dysfunction in disease progression is well-established, controversial findings concerning mt-dynamics have been described in MASLD animal models and minor evidence is reported in humans. Moreover, the impact of genetics on mt-lifecycle has been poorly investigated. Recently, we demonstrated that *PNPLA3*, *MBOAT7* and *TM6SF2* haploinsufficiency impair mt-biogenesis in hepatocytes through *PGC-1α*, the master regulator of mt-dynamics, causing an aberrant accumulation of warped mitochondria. Still, this model runs into mt-metabolic reprogramming, showing a high proliferative and invasive profile, thus corroborating the observations in humans (3).

Several studies underlined the urgent need to develop MASLD risk predictive models attempting to monitor disease progression and to bypass liver biopsy, the gold standard for MASLD diagnosis. Evaluation of mtDNA copy number (CN) and cell-free circulating mtDNA (ccf-mtDNA) are gaining attention for MASLD non-invasive assessment. mtDNA-CN may reflect mt-content and mt-dynamics, while ccf-mtDNA may be indicative of mt-functional status, as its release may be exacerbated by mt-damage which breaks mtDNA (11). Furthermore, the efforts for the development of reliable risk scores to apply in clinical practice is limited by the low accuracy rates observed (2, 12, 13). Nonetheless, the current advent in the use of artificial intelligence (AI) in medicine may pave the way for big data mining also in the absence of high-tech resources, thus allowing researchers to apply AI in different fields, from medical imaging to clinical data analysis.

To lend robustness to our *in vitro* data, aims of this study were to assess hepatic mt-dynamics in a Discovery cohort (n=28) and in a Validation cohort (n=824) of MASLD patients undergoing liver biopsy for diagnostic reasons and genetic screening stratified according to number of risk variants (NRV), as previously described (3). For the Discovery cohort, liver biopsies underwent a dedicated protocol for transmission electron microscopy (TEM) and an expanded panel of immunohistochemistry (IHC) analysis for the evaluation of mt-morphology and mt-dynamics markers, respectively. For the Validation cohort, the hepatic assessment of mt-biogenesis, content and damage was performed in a subset of liver biopsies (n=201). Mt-biomass was even estimated in MASLD-HCC specimens (n=7) and compared between intra-and extra-hepatic tissues and across genetic background. For both entire cohorts, we collected peripheral blood cells (PBMCs) and serum samples for the isolation of mtDNA-CN and ccf-mtDNA, respectively. To test their clinical utility, predictive non-invasive scores, including mitochondrial, anthropometric and genetic data, were developed by employing GPT-4 as AI statistical support and tested in the Validation cohort as potential diagnostic/prognostic tools for both early and late-stages of MASLD.

## Results

### *PNPLA3*, *MBOAT7* and *TM6SF2* SNPs exacerbate alterations of mt-morphology and content in the Discovery cohort

A case series of 28 newly enrolled biopsied MASLD patients (Discovery cohort; **Supplementary File** and **Table S1**) was included in the study to explore mt-morphological changes by TEM and to evaluate mt-dynamics by IHC according to both disease severity and genetic background. The assessment of mt-content and structure across MASLD histological damage (**Fig. S1A-D**) is described in the **Supplementary File.**

At the time of diagnosis, blood samples were collected for genotyping which identified only one non-carrier subject (3.5%; NRV=0), 11/28 (39.3%; NRV=1) carriers of one at-risk variant, 12/28 (42.8%; NRV=2) carriers of two mutations and 4/28 (14.3%; NRV=3) carrying all the three at risk polymorphisms (Table S1). NRV=3 patients showed the highest disease activity (NAS=6-7) and fibrosis grade (Fig. S2A-B), thus supporting that prevalence of disease severity progressively increases as the number of risk variants.

At TEM, we observed that NRV=3 patients displayed the highest mt-number (**Fig. 1A-D**; p=0.0001 at ANOVA; p=adj p=0.0004 *vs* NRV=0-1; adj p=0.008 *vs* NRV=2). At the generalized linear model (GLM), the hepatic mt-content inferred from TEM images, was associated with the presence of the three at-risk variants, independently of MASLD severity (**Table 1**), thus suggesting that genetics may directly alter mt-biogenesis.

**Fig. 1:**
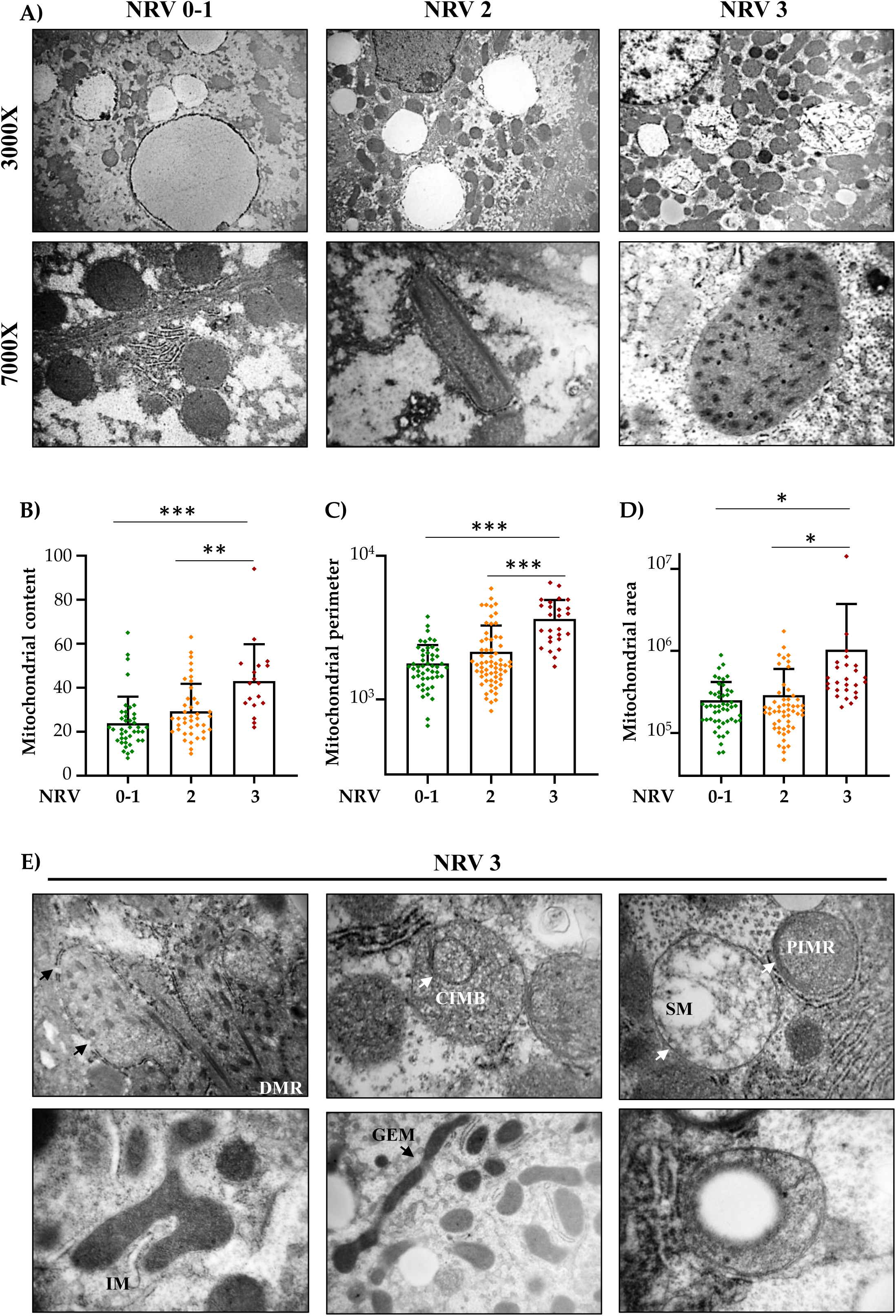
The effects of PNPLA3, MBOAT7 and TM6SF2 genetic variants on mt-morphology in the Discovery cohort. **A**) Representative TEM images showing mt-mass (on top, 3000X magnification) and canonical mt-defects as PIs and GMs (on bottom; 7000X magnification) in MASLD patients stratified according to NRV. **B**) Mt-content was calculated by counting mitochondria in micrographs at 3000X magnification through ImageJ. **C-D**) Mt-perimeter and area were measured through ImageJ software by tracing the boundary of mitochondria at 7000X magnification. Each data point in the bar graphs represents the mt-number (**B**) or mt-size (**C-D**) measured by ImageJ software in a range of 4-6 random images per patient. **E**) Noncanonical mt-morphological damages in NRV=3 MASLD patients included: DBR (double- membrane rupture); CIMB (circular internal membranes’ distribution); SM (swollen mitochondria); PIMR (packed internal membranes rearrangements); GEM (giant elongated mitochondria); VM (vacuolated mitochondria).

**Table 1.** Generalized linear model (GLM) correlating hepatic mt.-content and circulating *D-loop* expression measured by TEM and in PBMCs, respectively, with NAS≥5 and presence of NRV=3 variants in the Discovery cohort (n=28)

|  | Hepatic mitochondrial content |  | Circulating *D-loop* |  |
| --- | --- | --- | --- | --- |
| | $\beta$ {95% CI} | P value | $\beta$ {95% CI} | P value |
| Age, years | -0.19 {-0.54-0.15} | 0.25 | -0.03{-0.08-0.02} | 0.19 |
| BMI, kg/m<sup>2</sup> | -0.26 {-0.97-0.44} | 0.44 | -0.02{-0.11-0.07} | 0.67 |
| NAS (vs Mild) |  |  |  |  |
| ▪ Moderate <5 | -3.04 {-7.92-1.83} | 0.21 | 0.39{-0.28-1.06} | 0.24 |
| ▪ Severe ≥5 | 7.99 {3.98-11.99} | **0.0003** | 0.08{-0.47-0.64} | 0.73 |
| NRV=3 | 7.18 {3.37-10.99} | **0.0006** | 0.61{0.08-1.14} | **0.02** |

Values are reported as mean ± SD, number (%) or median {IQR}, as appropriate. BMI: body mass index; NAS: MASLD activity score; NRV: number of risk variants. GLM was adjusted for age, BMI, MASLD severity and NRV. Variables with skewed distribution (*D-loop* expression) were logarithmically transformed before analyses. p<0.05 was considered statistically significant.

Furthermore, NRV=3 patients presented the largest mt-size compared to those with 0, 1 or 2 variants (**Fig. 1C-D**; perimeter: p<0.0001 at ANOVA; adj p<0.0001 *vs* NRV=0-1 and NRV=2; area: p=0.02 at ANOVA; adj p=0.01 *vs* NRV=0-1 and NRV=2). Surprisingly, NRV=3 carriers accumulated a wide array of mt-morphological alterations including canonical defects (paracrystalline inclusions, PIs; giant mitochondria, GM), shared among subjects with a moderate/severe NAS (Supplementary File, Fig. S1A-D) and new ones as: a) rupture of mt-double membranes; b) abnormal distribution of the internal cristae (round-shaped; flattened sideways); c) swollen mitochondria with loss of internal matrix; d) irregular mt-morphology (three-branched); e) vacuoles inside rounded mitochondria (**Fig. 1E**).

These results suggest that the cumulative presence of *PNPLA3, TM6SF2* and *MBOAT7* SNPs exacerbate the hepatic mt- biogenesis and dysfunction, as shown by the increased mt-content and by the large spectrum of mt-ultrastructural lesions.

### *PNPLA3, MBOAT7* and *TM6SF2* mutations associated with hepatic PGC1a overexpression and unbalanced mt-dynamics in the Discovery cohort

Attempting to validate our hypothesis, we explored the hepatic expression of PGC-1α, the main transcriptional factor activating mt-biogenesis, and of downstream mediators regulating mt-lifecycle (fusion, fission and mitophagy) by IHC. PGC-1α was highly expressed in the hepatic parenchyma of NRV=3 MASLD patients (**Fig. 2A**). Specifically, NRV=0-1 patients showed ⁓20% of nuclear PGC-1α expression, and the amount of positive (+ve) cells increases of around 40% and 80% in NRV=2 and NRV=3 patients, respectively (p=0.001 at ANOVA; adj p=0.001 NRV=3 *vs* NRV=0-1; adj p=0.04 NRV=3 *vs* NRV=2; adj p=0.04 NRV=2 *vs* NRV=0-1, **Fig. 2B**, top), thus supporting that its transcriptional activity proportionally increases as the NRV. To give strength to our data, we marked mitochondria with a specific antibody recognizing mt-surface. As shown in **Fig. 2A** (bottom), mt-content significantly increased in NRV=3 patients (p=0.01 at ANOVA; adj p=0.02; NRV=3 *vs* NRV=0-1; **Fig. 2C**), forming large spotty marks, some of which close to the lipid droplets (LDs). Such observations confirmed the results obtained by TEM, suggesting that hepatic mt-mass increases in MASLD patients carrying the *PNPLA3*, *MBOAT7* and *TM6SF2* genetic mutations.

**Fig. 2:**
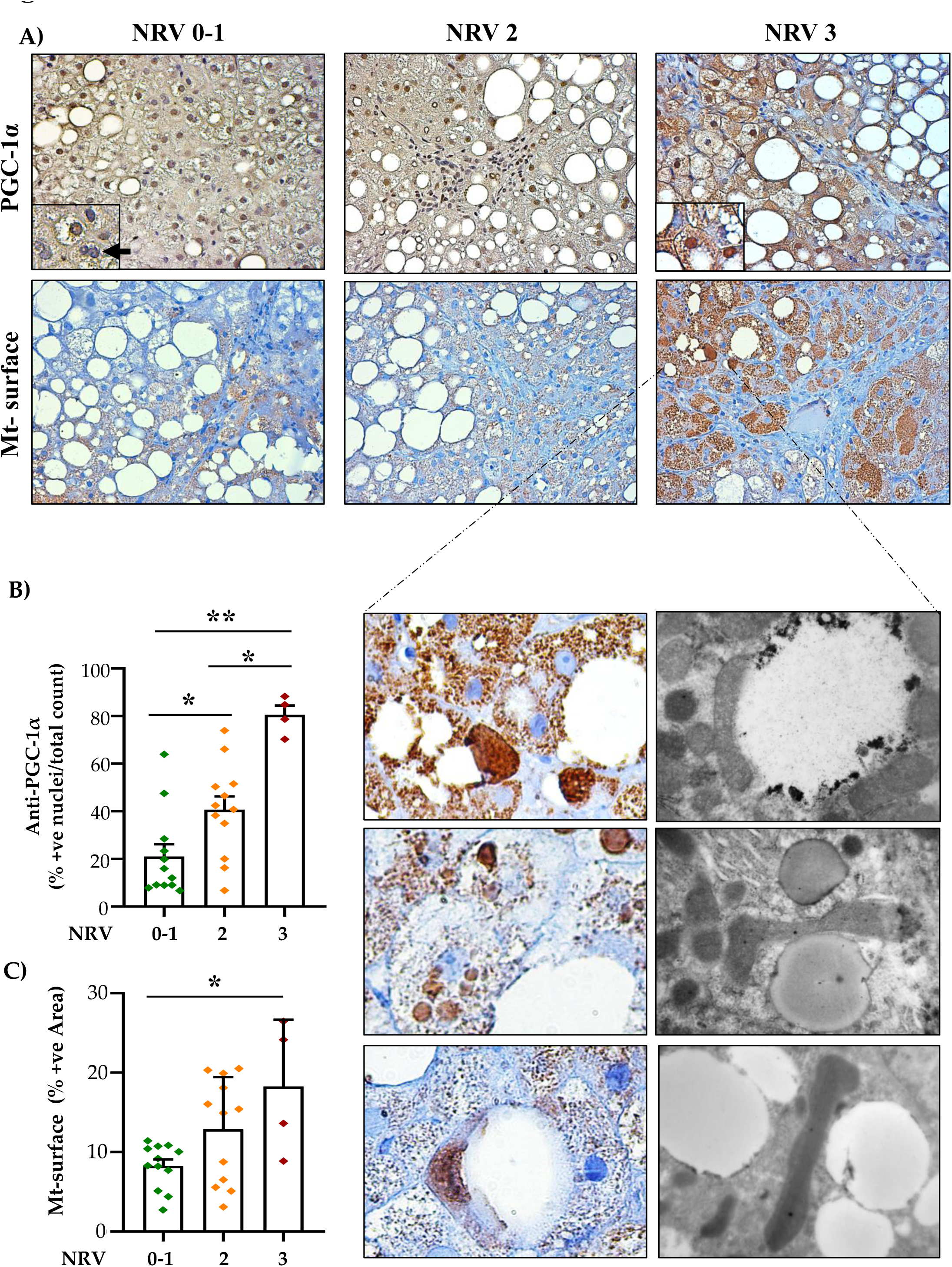
PNPLA3, MBOAT7 and TM6SF2 increases mt-mass in the Discovery cohort. **A**) Representative IHC images showing PGC-1α expression (on top, 40X magnification) and mitochondria (on bottom; 40X magnification) in MASLD patients stratified according to the NRV. **B**) Percentage of PGC-1α positive (+ve) cells was calculated by counting with ImageJ software the number of hepatocytes with brown-colored nuclei and normalizing data on total count of hepatocytes. **C**) Measurement of the area fraction per image, representing the positivity to the anti-mitochondrial surface antibody (brown-colored cytoplasm), was performed by splitting the RGB channels in red, green and blue separate components, setting a threshold and quantifying the brown intensity through ImageJ software. Each data point in the bar graphs represents the mean value of measured IHC images in a range of 4-6 random non-overlapping photos *per* patient.

Then, we moved on with IHC to explore markers involved in mt-cycle. The expression of Mitofusin 1 and 2 (MFN1/2), mediating aggregation of mt-outer membranes, was unchanged among MASLD patients stratified according to NRV (**Fig. 3A**). Conversely, mt-dynamin like GTPase (OPA1), involved in the same pathway of MFN1/2 but regulating mt-inner membranes fusion, was markedly increased in NRV=3 patients, capping around the LDs (p=0.005 at ANOVA; adj p=0.003 and adj p=0.04, NRV=3 *vs* NRV=0-1 or NRV=2; **Fig. 3A**, **Fig. S2C**), thereby indicating that it may play a key role in enhancing mt-fusion, potentially contributing to the formation of GM or in the irregular distribution of the inner mt-cristae detected by TEM.

**Fig. 3:**
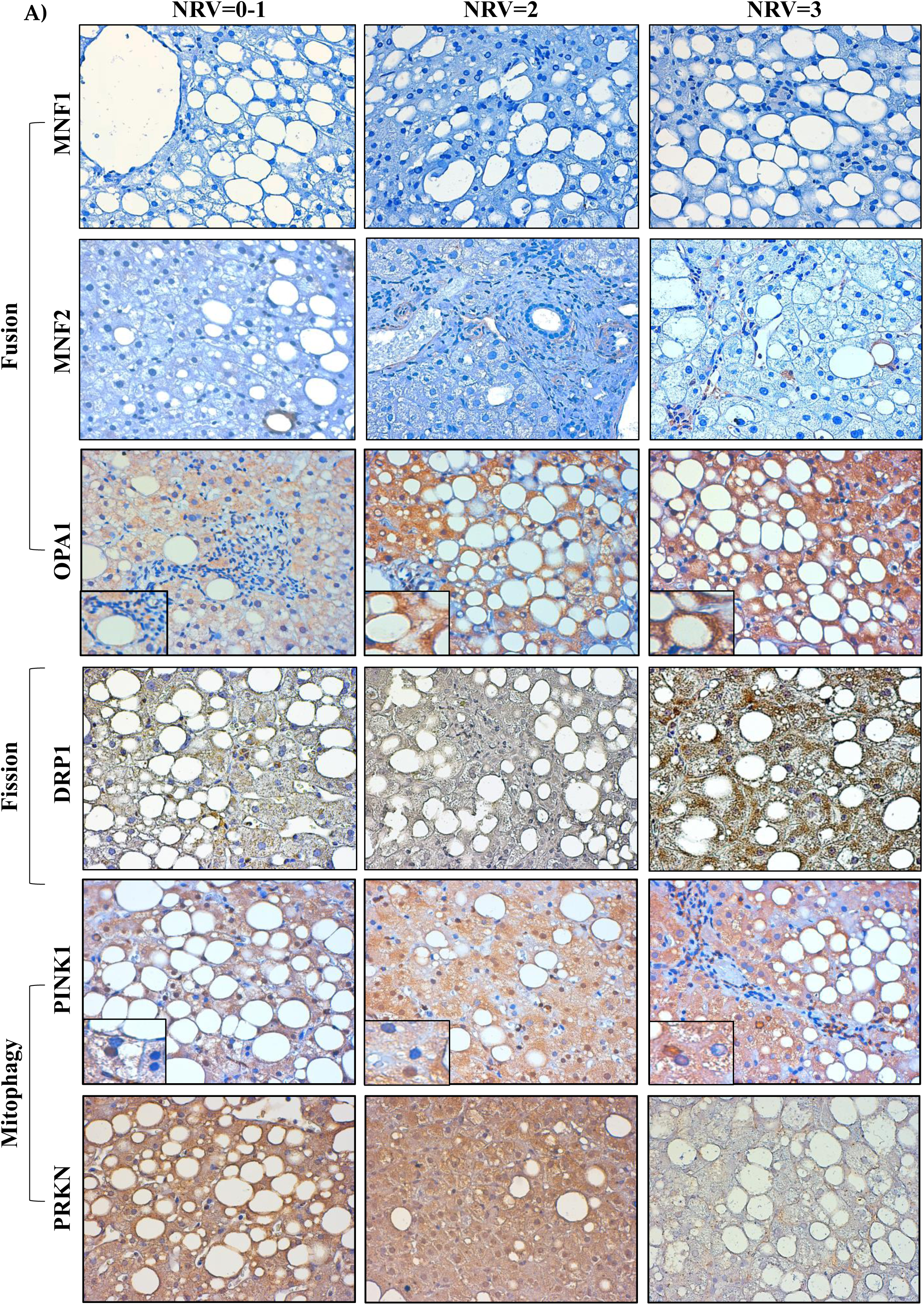
Characterization of hepatic mt-biogenesis in NRV=3 patients belonging to the Discovery cohort. **A**) A representative panel showing the expression of fusion proteins (MFN1/MFN2/OPA1), fission (DRP1) and mitophagy (PINK1, PRKN) assessed by IHC in liver biopsies of MASLD patients stratified according to NRV (40X magnification).

Still, dynamin-related protein 1 (DRP1) levels, promoting mt-fission, were higher in NRV=3 MASLD individuals compared to noncarriers or carriers of 1-2 variants (p=0.007 at ANOVA; adj p=0.0018; NRV=3 *vs* NRV=0-1 and NRV=2; **Fig. 3A**, **Fig. S2D**) and its expression strongly intensified in areas closest to the LDs. During mt-fission, mitophagy may be activated attempting to eliminate damaged mitochondria. Therefore, we finally investigated the expression of PINK1 and its downstream effector, Parkin (PRKN), principal orchestrators of mitophagy. We found that PINK1 was highly expressed in our cohort, mostly upregulated in the cytoplasm of hepatocytes belonging to patients with 2 variants and, even more, in those with NRV=3 (p=0.0002 at ANOVA, adj p=0.0006 vs NRV=0-1, **Fig. 3A**, **Fig. S2E**). Notably, PRKN expression was robust in patients harboring 0-1 or 2 variants, but it was markedly reduced in those with 3 mutations (p=0.02 at ANOVA; adj p=0.01 and adj p=0.04 vs NRV=0-1 or NRV=2, respectively, **Fig. 3A**, **Fig. S2F**). This observation suggests that while mitophagy is initiated in NRV=3 patients, the process may be early arrested, leading to the accumulation of non-functional mitochondria.

Collectively, these findings corroborated with the hypothesis that mt-biogenesis is augmented in NRV=3 carriers, as evidenced by a significant induction of OPA1 and DRP1. These proteins, in turn, may facilitate the formation of GM and small, round-shaped mitochondria, respectively, thereby contributing to the observed increase in mt-content. However, our findings have also revealed a plethora of ultrastructural abnormalities among the accumulated mitochondria in NRV=3 patients. These defects are likely resistant to complete eradication through the activation of the PINK1-PRKN pathway, given that the signaling cascade seems to be prematurely halted in these individuals.

### Dissecting the role of PNPLA3, MBOAT7 and TM6SF2 SNPs on mt-dynamics in the Discovery cohort

Then, we tried to elucidate the individual contributions of *PNPLA3, MBOAT7* and *TM6SF2* SNPs on mt-dynamics by analyzing the IHC images. We observed a notable increase in mt-biomass in individuals carrying either the *MBOAT7* rs641738 C>T variant or the TM6SF2 E167K polymorphism compared to those without these variants (p=0.004 at ANOVA; MBOAT7: adjusted p=0.02 for CC *vs* CT/TT; TM6SF2: adjusted p=0.001 for CC *vs* CT/TT; **Fig. 4A**). Conversely, hepatic mt-content unvaried among patients with or without the PNPLA3 at-risk genotype. These associations were substantiated by GLM, adjusted as above (**Table 2**), thus highlighting the critical roles of the *MBOAT7* and *TM6SF2* genes in the regulation of mt-mass.

**Fig. 4:**
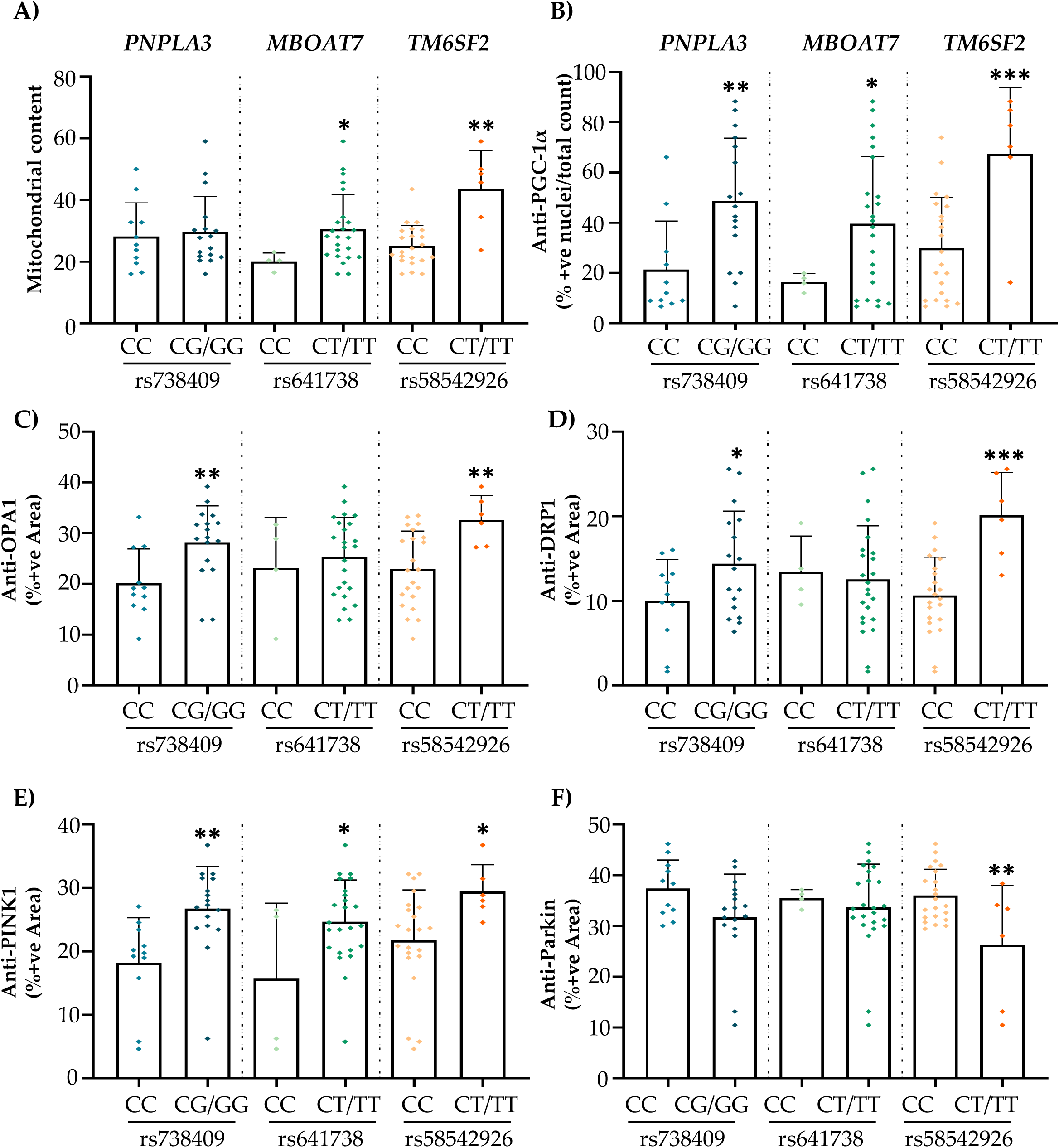
The individual contribution of PNPLA3, MBOAT7 and TM6SF2 genetic variants on both mt-mass and markers of mt-biogenesis in the Discovery cohort. **A**) Hepatic mitochondria were counted at 3000X magnification through ImageJ in MASLD patients stratified according to rs738409 C>G in *PNPLA3* (I148M), the rs641738 C>T in *MBOAT7-TMC4* locus and the rs58542926 C>T in *TM6SF2* (E167K) mutations. **B**) Percentage of PGC-1α positive (+ve) cells was calculated by counting with ImageJ software the number of hepatocytes with brown-colored nuclei and normalizing data on total count of hepatocytes. **C-F**) Measurement of the area fraction per image, representing the positivity of mt-markers (OPA1, DRP1, PINK1, PRKN; brown-colored cytoplasm), was performed by splitting the RGB channels in red, green and blue separate components, setting a threshold and quantifying the brown intensity through ImageJ software. Each data point in the bar graphs represents the mean value of measured IHC images (a range of 4-6 random non-overlapping photos *per* patient, 40X magnification).

**Table 2.** Generalized linear model (GLM) correlating hepatic mt-content measured by TEM with both NAS≥5 and presence of SNPs in *PNPLA3*, *MBOAT7-TMC4* and *TM6SF2* genes in the Discovery cohort (n=28)

| Hepatic mitochondrial content |  |  |
| --- | --- | --- |
| | $\beta$ {95% CI} | P value |
| Age, years | -0.27{-0.54-0.007} | 0.04 |
| BMI, kg/m<sup>2</sup> | -0.20{-0.72-0.31} | 0.41 |
| NAS (vs Mild) |  |  |
| ▪ Moderate <5 | -0.003{-3.82-3.81} | 0.99 |
| ▪ Severe ≥5 | 8.05{5.07-11.03} | **<0.0001** |
| *PNPLA3* G, yes | -0.07 {-2.16-2.00} | 0.93 |
| *MBOAT7* T, yes | 5.07 {1.98-2-8.15} | **0.002** |
| *TM6SF2* T, yes | 7.23{4.68-9.78} | **<0.0001** |

Values are reported as mean ± SD, number (%) or median {IQR}, as appropriate. BMI: body mass index; NAS: MASLD activity score; NRV: number of risk variants; SNPs: single nucleotide polymorphisms. GLM was adjusted for age, BMI, MASLD severity, the rs738409 C>G in *PNPLA3* (I148M), the rs641738 C>T in *MBOAT7-TMC4* locus and the rs58542926 C>T in *TM6SF2* (E167K). p<0.05 was considered statistically significant.

Immunohistochemical assessments of mt-markers provided additional insights about the role of *PNPLA3, MBOAT7* and *TM6SF2* mutations on mt-turnover. The prevalence of nuclear PGC-1α +ve hepatocytes was significantly higher in I148M PNPLA3 carriers compared to noncarriers (p=0.0005 at ANOVA; adjusted p=0.009 for CC *vs* CG/GG; **Fig. 4B**). This upregulation was consistent with the increase in OPA1, DRP1 and PINK1 +ve areas in individuals with *PNPLA3* CG/GG genotype (OPA1: p=0.01 at ANOVA; adj p=0.01 *vs* CC; DRP1: p=0.005 at ANOVA, adj p=0.04 *vs* CC; PINK1: p=0.002 at ANOVA, adj p=0.009 *vs* CC; **Fig. 4C-E**). Such observations supported the hypothesis that the *PNPLA3* rs738409 C>G mutation could stimulate mt-biogenesis in MASLD patients. Nonetheless, this did not correspond to an overall increase of mt-content, as one might anticipate (**Fig. 4A**, **Table 2**).

As concerns the *MBOAT7* rs641738 C>T variant, we found that T allele carriers exhibited an elevated nuclear PGC-1α positivity compared to noncarriers (p=0.003 at ANOVA, adj p=0.02 *vs* CC, **Fig. 4B**). However, this did not translate into significant alterations in the expression of mt-fusion or fission markers (**Fig. 4C-D**). Intriguingly, an augmented PINK1 +ve area fraction was observed, suggesting that the *MBOAT7* variant may influence mt-biomass through mitophagy modulation .

Interestingly, TM6SF2 E167K carriers showed the highest levels of PGC-1α +ve nuclei (p=0.0005 at ANOVA; adj p=0.002 CC *vs* CT/TT; **Fig. 4B**). This variant also correlated with upregulation of OPA1, DRP1, and PINK1 (OPA1: p=0.01 at ANOVA; adj p=0.01 *vs* CC; DRP1: p=0.005 at ANOVA, adj p=0.002 *vs* CC; PINK1: p=0.002 at ANOVA, adj p=0.04 *vs* CC; **Fig. 4C-E**), alongside a diminished PRKN +ve area fraction (p=0.04 at ANOVA, adj p=0.01 *vs* CC, **Fig. 4F**).

Taken together, these results indicated that the PNPLA3 I148M variant might contribute to an upregulation of mt-dynamics without affecting net mt-content, potentially through a compensatory balance in mt-turnover. Instead, the *MBOAT7* and *TM6SF2* mutations disrupt this equilibrium, promoting mt-biogenesis and impacting on mt-degradation, thus leading to significant alterations in mt-biomass.

### Non-invasive evaluation of mt-derived biomarkers in the Discovery cohort

Data obtained in liver biopsies revealed for the first time the relevant role of genetics in the modulation of mt-biogenesis. The next step was therefore to investigate whether this information could also be collected at circulating level by assessing biomarkers of mt-origin (**Supplementary File**).

In PBMCs, we measured *D-loop* levels, a noncoding region in the mtDNA replication site, which should proportionally reflect the amount of mtDNA-CN predicted from *D-loop* expression (**Supplementary file**) and the total mt-mass. Accordingly, we found that circulating *D-loop* expression positively correlated with hepatic mt-number quantified by TEM and with circulating mtDNA-CN (**Fig. S3A-B**). Considering the sample size of the cohort under investigation, both analyses showed meaningful correlation coefficients (r=0.387 and r=0.845, respectively), thereby supporting that *D-loop* may be a good candidate for the non-invasive evaluation of mt-mass. Moreover, *D-loop* levels were higher in NRV=3 patients compared to those with NRV=0-1 and NRV=2 (p=0.07 at ANOVA, adj p =0.01 *vs* NRV=0-1, adj p=0.04 *vs* NRV=2, **Fig. S3C**) and the association was significant at GLM adjusted for age, BMI and NAS (**Table 1**), thereby sustaining the results obtained by TEM/IHC and corroborating the hypothesis according to which the three at-risk genotypes act as independent modulators of mt-dynamics.

Instead, ccf-COXIII, a coding subunit of cytochrome c oxidase, whose fragment release may be commensurate to mt- damage, was chosen as serum parameter of mt-dysfunction. In NRV=3 patients, we found a heterogeneous range of modifications of the mt-structure, suggestive of an altered mt-activity. In accordance with TEM, serum ccf-COXIII levels were significantly higher in NRV=3 subjects compared to noncarriers or carriers of 1-2 variants (p=0.06 at ANOVA, adj p=0.03 *vs* NRV=0-1, adj p=0.02 *vs* NRV=2, **Fig. S3D**). However, no significant association emerged between ccf-COXIII and NRV=3 at multivariate analysis adjusted as above (β=0.03, 95% CI: -0.47-0.40, p=0.87), possibly ascribable to lack of statistical power.

### Hepatic assessment of mt-biomass in a subset of liver biopsies (n=201) from the Validation cohort

We then widened the characterization about mt-dynamics and its relationship with genetics in a larger cohort of 824 biopsied MASLD subjects (Validation cohort, **Supplementary file** and **Table S2**).

For the hepatic assessment of mt-dynamics and mt-biomass, we assessed gene expression analysis of *PGC-1α* and *D-loop* which were selected as representative markers, in a subset (n=201) of liver biopsies from the Validation cohort (**Supplementary File**). *PGC-1α* mRNA levels significantly increased in NRV=3 MASLD patients compared to NRV=0- 1 or NRV=2 subjects (p=0.06 at ANOVA, adj p=0.01 *vs* NRV=0-1; adj p=0.03 *vs* NRV=2, **Fig. S4A**). Accordingly, mtDNA-CN and *D-loop* levels were higher in liver biopsies of NRV=3 individuals rather than NRV=0-1-2 patients (mtDNA-CN: p=0.006 at ANOVA, adj p=0.001 *vs* NRV=0-1; adj p=0.04 *vs* NRV=2; *D-loop*: p=0.001 at ANOVA, adj p=0.02 *vs* NRV=0-1; adj p=0.03 *vs* NRV=2; **Fig. S4B-C**). As concerns mt-functions, we evaluated the hepatic expression of complexes of the oxidative phosphorylation (OXPHOS) and we found that UQCRC2, a subunit of the complex III, was significantly lower in NRV=3 patients compared to noncarriers or carriers of 1-2 variants (**Fig. S4D-E**).

Moreover, we had the availability to carry out a preliminary estimate of mt-biomass in n=7 hepatic tissues of MASLD-HCC patients, who underwent hepatic resection and of whom 4 had intra/extra tumoral counterparts. In this subset, three samples carried NRV=1, one patient had NRV=2 and three were NRV=3. The latter showed the highest mtDNA-CN and *D-loop* expression (**Fig. S4F-G**). In pair-matched MASLD-HCC tissues (n=4, intra- and extra-cancerous parts), a marked increase of mtDNA-CN was observed in tumoral lesions compared to extra-tumoral side (**Fig. S4H**) and the largest effect was found in a patient with NRV=3 (**Fig. S4I**).

Overall, these findings corroborated the results obtained in the Discovery cohort, providing further support for the substantial increase in mt-biomass and dysfunction in patients with a strong genetic predisposition and a greater disease severity. Notably, the risk of developing advanced disease is further accentuated in NRV=3 subjects to the extent that the co-existence of the three SNPs and MASLD-HCC results in the highest mt-content.

### Non-invasive evaluation of mt-derived biomarkers in the Validation cohort: the role of genetic background

In the Discovery cohort, we observed a positive association among circulating *D-loop* levels, ccf-COXIII and NRV=3 variants, supporting the hypothesis that these biomarkers could be surrogates for assessing alterations of mt-mass and functions in the bloodstream. These findings were strengthened in the entire Validation cohort (n=824 patients, of whom 125 MASLD-HCC) by both bivariate analysis (*D-loop*: p=0.04 at ANOVA, adj p=0.02 *vs* NRV=0-1; adj p=0.04 *vs* NRV=2, ccf-COXIII: p<0.0001 at ANOVA, p<0.0001 *vs* NRV=0-1-2, **Fig. 5A-B**) and GLM adjusted as above (*D-loop*: β=0.17, 95% CI: 0.04-0.29, p=0.008; ccf-COXIII: β=0.35, 95% CI: 0.22-0.48, p<0.0001; **Table 3**), thus corroborating the direct involvement of genetics in mt-dynamics and function.

**Fig. 5:**
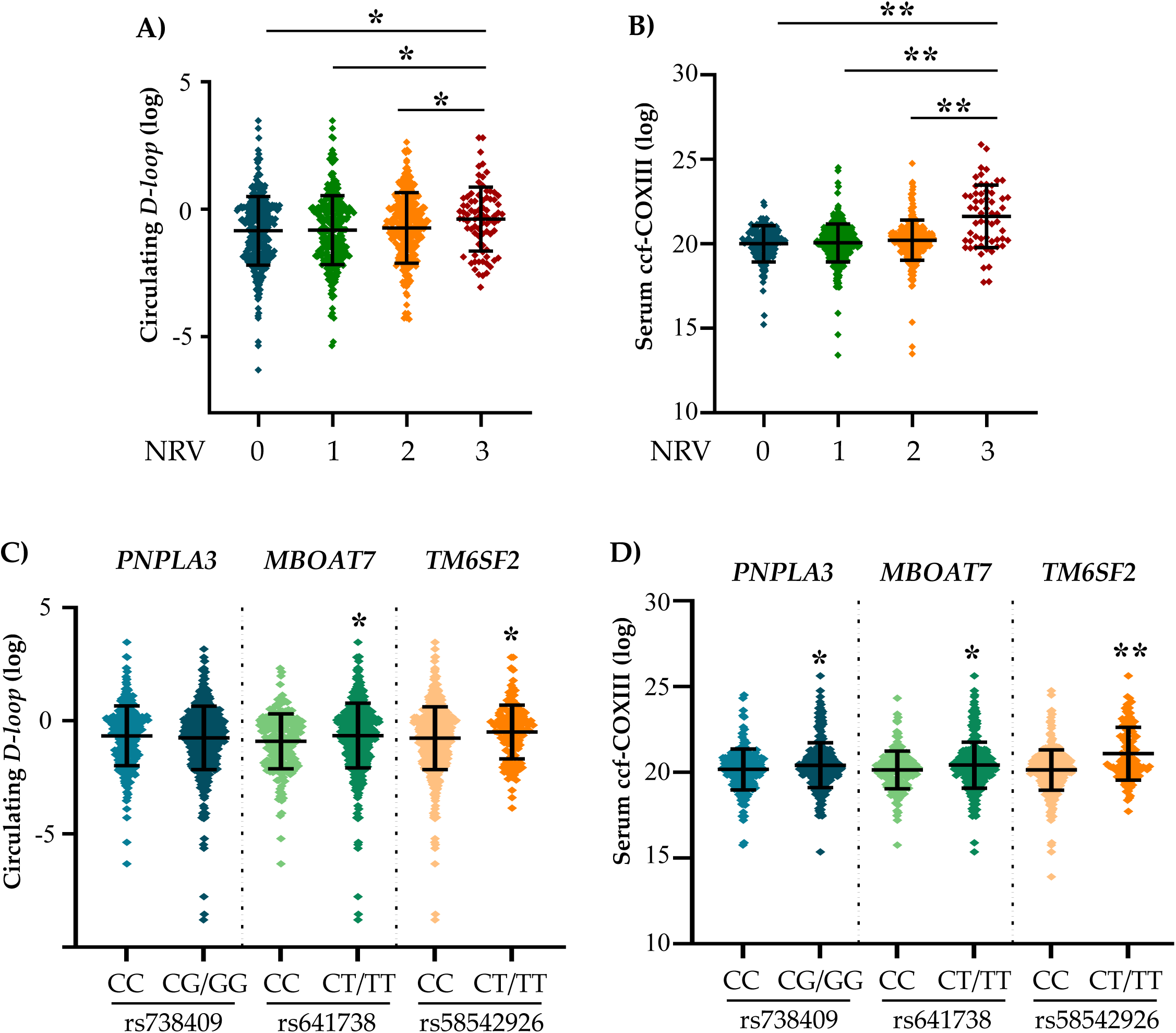
Evaluation of mt-derived biomarkers in MASLD patients of the Validation cohort (n=824). **A-B**) Bivariate analysis showing the cumulative weight of NRV=3 SNPs on circulating *D-loop* and serum ccf-COXIII levels across MASLD patients. **C-D**) Individual contribution of *PNPLA3* (I148M), the rs641738 C>T in *MBOAT7-TMC4* and *TM6SF2* (E167K) variants on changes in circulating *D-loop* and serum ccf-COXIII levels in the Validation cohort.

**Table 3.** On top, generalized linear mode (GLM)l correlating circulating *D-loop* expression measured in PBMCs and ccf- COXIII serum levels, with the presence of NRV=3 variants and NAS in the Validation cohort (n=824). On bottom, GLM correlating both biomarkers with presence and presence of SNPs in *PNPLA3*, *MBOAT7-TMC4* and *TM6SF2* genes.

| Circulating *D-loop* |  |  | Serum ccf-COXIII |  |
| --- | --- | --- | --- | --- |
| | $\beta$ {95% CI} | P value | $\beta$ {95% CI} | P value |
| Age, years | 0.002 {-0.005-0.01} | 0.47 | 0.0007 {-0.007-0.008} | 0.85 |
| BMI, kg/m<sup>2</sup> | -0.003 {-0.016-0.02} | 0.74 | -0.014 {-0.03-0.008} | 0.21 |
| NAS (vs Mild) |  |  |  |  |
| ▪ Moderate <5 | 0.05 {-0.08-0.29} | 0.45 | 0.06 {-0.07-0.20} | 0.36 |
| ▪ Severe $\geq 5$ | 0.21 {0.006-0.36} | **0.005** | 0.05 {-0.20-0.09} | 0.48 |
| NRV=3 | 0.17 {0.04-0.29} | **0.0006** | 0.35 {0.22-0.48} | **<0.0001** |
| | $\beta$ {95% CI} | P value | $\beta$ {95% CI} | P value |
| Age, years | 0.002 {-0.005-0.01} | 0.57 | 0.001 {-0.006-0.009} | 0.72 |
| BMI, kg/m<sup>2</sup> | -0.004 {-0.015-0.02} | 0.66 | -0.13 {0.03-0.009} | 0.24 |
| NAS (vs Mild) |  |  |  |  |
| ▪ Moderate <5 | 0.07 {-0.07-0.21} | 0.33 | 0.07 {-0.07-0.21} | 0.31 |
| ▪ Severe $\geq 5$ | 0.22 {0.07-0.37} | **0.003** | 0.08 {-0.07-0.23} | 0.29 |
| *PNPLA3* G, yes | 0.04 {-0.05-0.14} | 0.40 | 0.13 {0.02-0.23} | **0.01** |
| *MBOAT7* T, yes | 0.17 {0.07-0.27} | **0.001** | 0.11 {0.01-0.22} | **0.02** |
| *TM6SF2* T, yes | 0.13 {0.003-0.25} | **0.04** | 0.35 {0.21-0.48} | **<0.0001** |

Values are reported as mean ± SD, number (%) or median {IQR}, as appropriate. BMI: body mass index; NAS: MASLD activity score; NRV: number of risk variants. The multivariate was adjusted for age, BMI, NAS and presence of rs738409 C>G in *PNPLA3* (I148M), the rs641738 C>T in *MBOAT7-TMC4* locus and the rs58542926 C>T in *TM6SF2* (E167K) alone or combined (NRV). p<0.05 was considered statistically significant.

In accordance with the prior approach, we carried out a sub-analysis attempting to assess the individual contribution of PNPLA3 I148M, *MBOAT7* rs641738 C>T and TM6SF2 E167K mutations in modulating circulating *D-loop* and ccf- COXIII levels. As concerns the former, at bivariate analysis, carriers of both *MBOAT7* rs641738 C>T and TM6SF2 E167K variants showed elevated *D-loop* expression compared to noncarriers (p=0.03 at ANOVA; MBOAT7: adj p=0.014 CT/TT *vs* CC; TM6SF2: adj p=0.015 CT/TT *vs* CC, **Fig. 5C**), while no effects were detected with the PNPLA3 I148M SNP. These associations remained significant at GLM adjusted for confounding variables such as age, BMI and NAS and presence of PNPLA3 I148M variant (*MBOAT7*: β=0.17, 95% CI: 0.07-0.27, p=0.001; *TM6SF2*: β=0.13, 95% CI: 0.003-

0.25, p=0.04, **Table 3**), thus reinforcing the observations of the Discovery cohort regarding the contribution of individual variants on changes in mt-content. Indeed, these findings indicate that the *PNPLA3* genetic mutation has a lesser impact on circulating marker, which should non-invasively reflect mt-mass, compared to *MBOAT7* and *TM6SF2* which, on the contrary, appear to have a predominant role on this aspect.

Serum ccf-COXIII increased in carriers of PNPLA3 I148M, *MBOAT7* rs641738 C>T and TM6SF2 E167K SNPs (p<0.001 at ANOVA; PNPLA3: adj p=0.01 CG/GG *vs* CC; MBOAT7: adj p=0.02 CT/TT *vs* CC; TM6SF2: adj p<0.0001 CT/TT *vs* CC, **Fig. 5D**) and it correlated with all three single mutations at multivariate analysis adjusted as before (*PNPLA3*: β=0.13, 95% CI: 0.02-0.23, p=0.01; *MBOAT7*: β=0.11, 95% CI: 0.01-0.22, p=0.02; *TM6SF2*: β=0.35, 95% CI: 0.21-0.48, p<0.0001, **Table 3**).

Therefore, data obtained in the Validation cohort clearly highlighting that *MBOAT7* and *TM6SF2* genetic mutations, but not the PNPLA3 I148M variant, play a significant role in altering total mt-number, while the effects on mt-functions seems to be independently affected by all three SNPs.

### Performance of mt-derived biomarkers in clinical risk assessment

To investigate the potential clinical utility of mt-derived biomarkers, we correlated both circulating *D-loop* and ccf- COXIII levels with anthropometric and histological data of the Validation cohort. With the goal to develop and test novel risk scores based on the ability of both biomarkers in identifying both early and late forms of MASLD, the Validation cohort was stratified according the to the histological damage as follows: SLD, MASH, fibrosis and MASLD-HCC (**Table S2**).

### D-loop: a pathologic sign of MASLD progression

A positive association between *D-loop* expression and BMI, T2DM, HOMA-IR, AST, lactate and LDH emerged at bivariate (**Fig. S5A-F**) and multivariate analyses, adjusted for confounding factors implicated in disease progression (sex, age, BMI, T2DM and NRV=3), (**Table S3**), suggesting that mt-mass changes in response to MASLD-related metabolic comorbidities and increases with indices of liver damage.

Accordingly, at bivariate analysis *D-loop* expression progressively increased with disease severity (SLD: 0.31{0.12- 0.94}; MASH: 0.44{0.19-0.96}, fibrosis: 0.57{0.18-1.14}; MASLD-HCC: 0.82{0.46-1.46}; p=0.01; **Table S2**) and its levels correlated with severe NAS at GLM (β=0.21, 95% CI: 0.006-0.36, p=0.005, **Table 3**), a relationship which was not previously detected in the Discovery cohort possibly due to the poor sample size.

These data were corroborated by the ordinal logistic regression analysis corrected as above, which revealed that *D-loop* was independently associated with the histological degree of steatosis (β=0.08, 95% CI: 0.01-0.15, p=0.02) necroinflammation (β=0.09, 95% CI: 0.21-0.17, p=0.01), ballooning (β=0.18, 95% CI: 0.09-0.27, p<0.0001) and fibrosis (β=0.08, 95% CI: 0.003-0.15, p=0.03) (**Table 4**). Still, at nominal logistic regression analysis *D-loop* levels correlated with increased risk of MASH, cirrhosis and MASLD-HCC (OR MASH: 1.27, 95% CI: 1.08-1.48, p=0.02; OR cirrhosis:1.32, 95% CI: 1.06-1.63, p=0.008; OR MASLD-HCC: 1.31, 95% CI: 1.01-1.71, p=0.03, **Table 5**). Taken together, such findings supported that higher *D-loop* levels correlate with environmental factors, genetics and MASLD severity, thus representing an independent risk factor of MASLD progression.

**Table 4.** Ordinal logistic regression analysis correlating circulating *D-loop* levels with histological damage in the Validation cohort (n=824)

| Steatosis |  |  | Necroinflammation |  |
| --- | --- | --- | --- | --- |
| | $\beta$ {95% CI} | *P* value | $\beta$ {95% CI} | *P* value |
| Sex, M | 0.27{0.15-0.39} | <0.0001 | 0.29{0.16-0.41} | <0.0001 |
| Age, years | 0.001{-0.007-0.01} | 0.72 | 0.02{0.01-0.03} | <0.0001 |
| BMI, kg/m<sup>2</sup> | 0.03{0.02-0.04} | <0.0001 | 0.02{0.006-0.03} | 0.0046 |
| IFG/T2D, yes | 0.32{0.19-0.46} | <0.0001 | 0.32{0.18-0.46} | <0.0001 |
| NRV=3 | 0.46{0.32-0.61} | <0.0001 | 0.27{0.13-0.42} | 0.0002 |
| *D-loop* (log) | 0.08{0.01-0.15} | **0.02** | 0.09{0.02-0.17} | **0.01** |
| Ballooning |  |  | Fibrosis |  |
| | $\beta$ {95% CI} | *P* value | $\beta$ {95% CI} | *P* value |
| Sex, M | 0.28{0.13-0.42} | 0.0001 | 0.34{0.22-0.46} | <0.0001 |
| Age, years | 0.03{0.02-0.04} | <0.0001 | 0.05{0.04-0.06} | <0.0001 |
| BMI, kg/m<sup>2</sup> | -0.009{-0.026-0.008} | 0.30 | -0.002{-0.01-0.012} | 0.74 |
| IFG/T2D, yes | 0.31{0.15-0.46} | <0.0001 | 0.57{0.43-0.71} | <0.0001 |
| NRV=3 | 0.30{0.12-0.48} | 0.0006 | 0.37{0.23-0.51} | <0.0001 |
| *D-loop* (log) | 0.18{0.09-0.27} | **<0.0001** | 0.07{0.003-0.15} | **0.03** |

Values are reported as mean ± SD, number (%) or median {IQR}, as appropriate. BMI: body mass index; IFG/T2D: impaired fasted glucose/ type 2 diabetes; NAS: MASLD activity score; NRV: number of risk variants. Beta ( $\beta$ ) ± SE (standard error); CI: confidence interval. Multivariate analysis was adjusted for age, BMI, NAS, the cumulative presence of rs738409 C>G in *PNPLA3* (I148M), the rs641738 C>T in *MBOAT7-TMC4* locus and the rs58542926 C>T in *TM6SF2* (E167K) and circulating *D-loop*, which was logarithmically transformed. $p < 0.05$ was considered statistically significant.

**Table 5.**
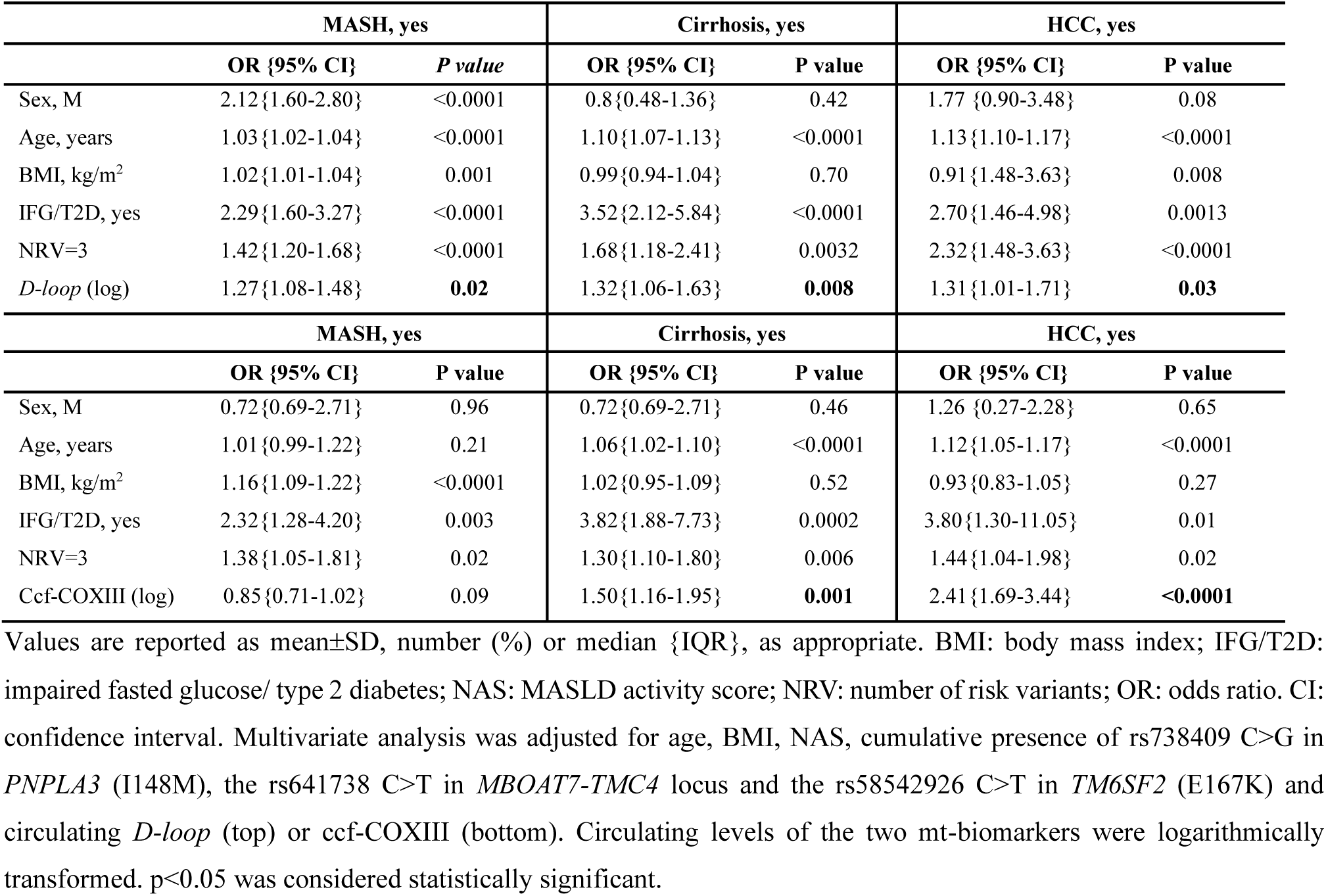
Nominal logistic regression analysis correlating circulating *D-loop* levels and serum ccf-COXIII with risk of MASH, cirrhosis and HCC in the Validation cohort (n=824)

### ccf-COXIII: a prognostic indicator of MASLD-HCC

Differently from *D-loop*, ccf-COXIII did not correlate with biochemical and anthropometric variables associated to MASLD and its serum quantity did not progressively increase across SLD, MASH and fibrosis stages (**Tables S2**- **S4**). In keeping with this evidence, at multivariate analysis adjusted for sex, age, BMI, T2DM and NRV=3, ccf-COXIII showed a trend of correlation with the histological parameters of liver damage and MASH (**Table S5, Table 5**), but the analysis did not reach the statistical significance.

Nonetheless, ccf-COXIII levels were dramatically higher in MASLD-HCC patients compared to those with SLD, MASH and fibrosis (**Table S2**). At nominal logistic regression analysis corrected as above, serum ccf-COXIII showed a significant association with the worst stages of the disease, increasing the risk of cirrhosis and MASLD-HCC by around 1.5-fold and 2.4-fold, respectively (**Table 5**). Therefore, MASLD patients with the highest disease severity present a massive release of ccf-COXIII fragment, supporting that its concentration proportionally fits with the intensity of hepatic injury and mt-dysfunction. Despite ccf-COXIII is less informative than *D-loop* for the risk of MASH, it emerged as a promising prognostic measure of advanced MASLD.

### Risk scores development through GPT-4: the role of mt-biomarkers as prognostic factors of MASLD-HCC

Our analyses revealed that the two mt-biomarkers may have different values: *D-loop* is likely influenced by comorbidities and increased with disease severity, while ccf-COXIII specifically rises in HCC cases. In both cases, their assessment may play a relevant part in clinical practice for the early identification and monitoring of MASLD patients at higher risk to progress towards advanced conditions.

With the final outcome to evaluate their clinical efficiency, we developed novel risk scores tailored to the varying MASLD forms: the MAGIC-MASH for detecting early-stage MASH, MAGIC-Fib for assessing fibrosis, and the MAGIC-H for identifying HCC. These scores were generated through AI-guided analysis using a customized version of GPT-4 (rsGPT- 4) according to the **Supplementary File** description.

After anonymizing the dataset, we uploaded it to the rsGPT-4 environment, which scanned the data to optimize the approach. We asked rsGPT-4 to develop the risk scores based solely on non-invasive biomarkers such as *D-loop* and ccf- COXIII, anthropometric and genetic data (age, BMI and NRV). Given our dataset characteristics, the AI identified a machine learning approach (Random Forest, RF), to obtain a high-performance risk score, which we manually validated on JMP software (SAS, Cary, NC). The RF analyses provided the relative feature importance of the tested parameters and of their interactions, which were subsequently gathered in formulas keeping into account the normalized, importance-weighted predictors (**Supplementary File**). We then applied the risk scores to the Validation cohort, and we generated AUC-ROC curves to determine their prognostic precision on the desired outcomes.

The MAGIC-MASH score built with *D-loop* alone showed a good accuracy (AUC: 73%) for MASH detection, with a discrete balance of sensitivity (65.7%) and specificity (71.3%). Conversely, the ccf-COXIII’s AUC was slightly lower at 67%, suggesting it is less discriminative than the *D-loop* for MASH. However, when both biomarkers were combined, the AUC was 69%, showing an increase in sensitivity (73.4%) at the expense of specificity (57.9%). This might suggest that combining these two biomarkers does not necessarily improve the model’s ability to discriminate between MASH and non-MASH cases compared to the *D-loop* alone (**Table 6**, **Fig. 6A**).

**Fig. 6:**
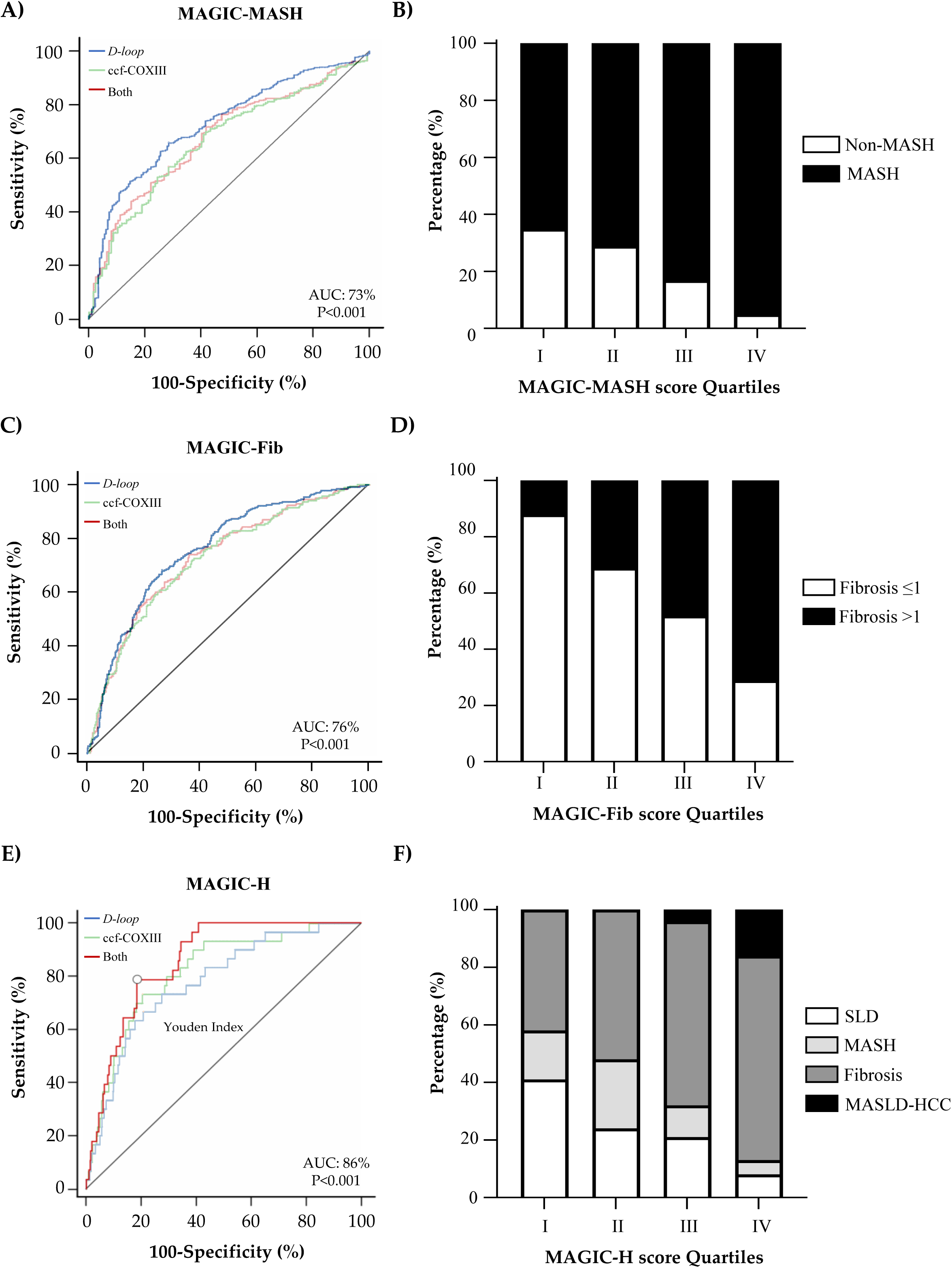
Prognostic accuracy of MAGIC- risk scores for MASLD clinical assessment, generated through rsGPT-4 guidance and random forest analysis in the Validation cohort. **A**) AUC-ROC curves comparing MAGIC-MASH accuracy, including mt-biomarkers alone or combined integrated with genetic background (NRV) and anthropometric parameters (age, BMI). **B**) MAGIC-MASH quartiles based on the best MAGIC-MASH score (calculated with *D-loop* alone) displaying the percentage (%) of MASH and non-MASH patients across classification. **C**) AUC-ROC curves comparing MAGIC-Fib accuracy, including mt-biomarkers alone or combined integrated with NRV, age, BMI. **D**) Contingency analysis showing the frequency of MASLD patients with fibrosis >1 falling into MAGIC-Fib quartiles, based on the best MAGIC-Fib score (calculated with *D-loop* alone). **E**) AUC-ROC curves comparing MAGIC-H accuracy, including mt- biomarkers alone or combined integrated with NRV, age, BMI. **D**) Contingency analysis shows the distribution (%) of MASLD patients, with different disease severity, falling into each MAGIC-H quartile. MedCalc software (MedCalc Software Ltd, Belgium) was used to extrapolate sensitivity, specificity and cut-offs of each score. Transparent lines depict alternative scoring models with lesser predictive power, whereas the marked lines highlight the most efficient score identified by the random forest analysis.

**Table 6.** AUC-ROC curves comparison performed by MedCalc software, which correlate rsGPT4-based MASLD risk score (MAGIC-) with MASH, fibrosis>1 and HCC risk in the Validation cohort (n=824).

| MASH, yes |  |  |  |
| --- | --- | --- | --- |
|  | *D-loop* | Ccf-COXIII | Both |
| AUC | 0.73 | 0.67 | 0.69 |
| 95% CI | 0.69-0.75 | 0.63-0.71 | 0.65-0.73 |
| Z statistics | 11.11 | 6.91 | 7.54 |
| Youden Index J | 0.40 [0.29-0.42] | 0.29 [0.18-0.38] | 0.31 [0.21-0.36] |
| Cut-off | >24.1 | >34.86 | >24.1 |
| Sensitivity (%) | 65.7 | 53.4 | 73.4 |
| Specificity (%) | 71.3 | 75.5 | 57.9 |
| **Sample size** | n=824 | n=824 | n=824 |
| ▪ Positive | 635 (77%) | 613 (74.4%) | 626 (77.8%) |
| ▪ Negative | 189 (22.8%) | 211 (26.6%) | 198 (24%) |
| Fibrosis>1, yes |  |  |  |
|  | *D-loop* | Ccf-COXIII | Both |
| AUC | 0.76 | 0.72 | 0.73 |
| 95% CI | 0.72-0.78 | 0.68-0.75 | 0.69-0.77 |
| Z statistics | 14.28 | 9.59 | 10.34 |
| Youden Index J | 0.41 [0.33-0.4] | 0.35 [0.26-0.40] | 0.38 [0.29-0.44] |
| Cut-off | >26.0 | >36.6 | >30 |
| Sensitivity (%) | 68.0 | 57.4 | 74.0 |
| Specificity (%) | 73.2 | 77.6 | 64.4 |
| **Sample size** | n=824 | n=824 | n=824 |
| ▪ Positive | 323 (39.2%) | 55 (6.7%) | 289 (35%) |
| ▪ Negative | 501 (60.8%) | 768 (93.3%) | 535 (64.9%) |
| HCC, yes |  |  |  |
|  | *D-loop* | Ccf-COXIII | Both |
| AUC | 0.77 | 0.81 | 0.86 |
| 95% CI | 0.74-0.81 | 0.78-0.85 | 0.82-0.88 |
| Z statistics | 14.19 | 9.71 | 17.17 |
| Youden Index J | 0.51 [0.39-0.58] | 0.55 [0.40-0.64] | 0.60 [0.54-0.66] |
| Cut-off | >28.9 | >38.3 | >30 |
| Sensitivity (%) | 73.5 | 75.7 | 78.6 |
| Specificity (%) | 77.5 | 80.1 | 81.5 |
| **Sample size** | n=824 | n=824 | n=824 |
| ▪ Positive | 92 (11.16%) | 55 (6.7%) | 44 (5.4%) |
| ▪ Negative | 732 (88.8%) | 768 (93.3%) | 780 (94.6%) |

AUC: area under the curve; CI: confidence interval. MAGIC- risk scores aggregated age, BMI, genetic background (PNPLA3, TM6SF2 and MBOAT7-TMC4 risk variants, NRV) and mt-biomarkers (*D-loop* and/or ccf-COXIII). MAGIC- formulas were achieved by random forest analysis, which weighted the relative importance of the abovementioned parameters (age, BMI, NRV, *D-loop*, and ccf-COXIII). Further, the interactions among *D-loop*/ccf-COXIII and NRV was considered in the analysis.

As increasing *D-loop* levels in concomitance with the presence of NRV=3 correlate with higher MAGIC-MASH score ranking, we moved forward to estimate the risk of developing MASH by stratifying patients according to MAGIC-MASH quartiles with the best AUC among those examined. Therefore, we classified subjects into quartiles of the MAGIC-MASH calculated using the single biomarker *D-loop* as follows: very low-risk (1^st^ quartile: ≤21.65 points), low-risk (2^nd^ quartile: 21.65 to <24.97 points), moderate-risk (3^rd^ quartile: 24.97 - <27.99 points), and high-risk (4^th^ quartile: ≥27.99 points). Consistently. we observed that patients belonging to the 4^th^ quartile had a significant increased risk of MASH compared to other quartiles (OR 9.89, 95% CI 5.05-19.37, p<0.0001 vs 1^st^ quartile; OR 7.44, 95% CI 3.66-15.12, p<0.0001 vs 2^nd^ quartile; OR 3.79, 95% CI 1.80-7.97, p=0.0004 vs 3^rd^ quartile) (**Fig. 6B**).

When we built the MAGIC-Fib score by considering *D-loop* and ccf-COXIII separately, we observed that *D-loop*, with an AUC of 76%, was again the strongest single biomarker for detecting fibrosis compared to ccf-COXIII (72%). When both biomarkers were included in the MAGIC-Fib, the AUC increases marginally to 73%. These values suggest good discriminative ability, though not substantially different when using both biomarkers versus the *D-loop* alone (**Table 6**, **Fig. 6C**). By applying a strategy similar to that used for MAGIC-MASH, we divided the cohort with MAGIC-FIB quartiles that considers the contribution of *D-loop* alone alongside the co-presence of NRV=3 to predict the risk of fibrosis: very low-risk (1^st^ quartile: ≤<21.83 points), low-risk (2^nd^ quartile: 21.83 to <25.43 points), moderate-risk (3^rd^ quartile: 25.43 - <28.69 points), and high-risk (4^th^ quartile: ≥28.69 points). This division yielded similar findings as above for MAGIC-MASH. In particular, patients in the 4^th^ quartile showed a proportional increased risk of fibrosis compared to patients in the other quartiles (OR 18.43, 95% CI 10.6-32.06, p<0.0001 vs 1^st^ quartile; OR 5.46, 95% CI 3.47-8.58, p<0.0001 vs 2^nd^ quartile; OR 2.56, 95% CI 1.65-3.96, p<0.0001 vs 3^rd^ quartile) (**Fig. 6D**).

Finally, the most notable increase in prognostic precision was observed in the MAGIC-H score for HCC risk stratification. Here, ccf-COXIII demonstrated a higher AUC than *D-loop* (81% *vs* 77%, respectively), thus sustaining that ccf-COXIII abundance is a stronger HCC predictor on its own. The dual-biomarkers model reached an AUC of 86% (95% CI 0.823- 0.885) with a sensitivity of 78.6% and specificity of 81.5%, thus optimizing the detection of HCC, particularly at a cut- off risk score set around 30 points (95% CI: 27.91-31.59, **Table 6**, **Fig. 6E**).

To further dissect the predictive power of the MAGIC-H score, we grouped the MASLD population into risk score quartiles including both mt-biomarkers (1^st^ quartile: <23.21 points; 2^nd^ quartile: 23.21 to <26.50 points; 3^rd^ quartile: 26.50 - <29.28 points; 4^th^ quartile: ≥29.28 points). At nominal logistic regression analysis, we observed that patients belonging to the 4^th^ quartile showed a significantly increased risk of HCC compared to 1^st^ (OR 24.37, 95% CI 4.22-184, p=0.002), 2^nd^ (OR 22.69, 95% CI 3-171, p=0.002) and 3^rd^ (OR 3.75, 95% CI 1.46-9.62, p=0.006) quartiles (**Fig. 6F**). Overall, the employment of rsGPT-4 has allowed to generate new risk models, whose high-level of accuracy is weighted on the relative contribution of demographic, genetic and mt-derived biomarkers, solidifying the models’ discriminative abilities.

## Discussion

The incidence of MASLD has exponentially spread worldwide, becoming one of the most troubling social and health alarm clocks (14). Predicted to exceed more than 50% by 2040, this global growth is certainly related to the increasing rates of obesity and diabetes (15). In parallel, the role of genetic traits has gained recognition for clinical risk assessment, highlighting the utility of polygenic risk scores (PRS) in identifying patients at high-risk of disease progression (2–6). Nonetheless, this aspect, not yet a routinary part of clinical practice, is often underscored or only partially addressed, thus also restricting the screening options for patients (16). Additionally, the development of PRS for clinical application faces challenges due to lack of precision, necessitating compromises in sensitivity or specificity that increase the risk of false positives or negatives (2, 12, 13).

Generative AI tools are increasingly used in healthcare to facilitate information gathering, foster discussions among physicians, patients, or relatives, and assist in digital pathology. However, the potential for AI in medical fields remains vast, particularly in enhancing diagnostic accuracy, treatments, and patient outcomes, many of which are not achieved yet (17, 18). Therefore, in the latest years, we focused our efforts on meeting the gap related to the impact of the major MASLD-associated genetic variants in *PNPLA3*, *TM6SF2* and *MBOAT7-TMC4* genes on disease worsening. Specifically, we attempted to address the following queries: 1) how these SNPs, involved in lipid remodeling, may lead to advanced MASLD forms in humans; 2) how to maximize the use of genetic variants in medical practice by proposing a novel analytical method under the AI support.

Previously, we demonstrated that a link exists between the co-presence of *PNPLA3*, *MBOAT7-TMC4* and *TM6SF2* loss-of-functions and the impairment of mt-flexibility in hepatocytes. We pointed out, for the first time, that *MBOAT7* and *TM6SF2* genes are crucial for maintaining the inter-organelles dynamic interplay and turnover in genetically-edited hepatocytes, possibly affecting membranes’ fluidity and lipid composition. Notably, the combined MBOAT7^-/-^TM6SF2^-/-^ model further showed a clear switching of mt-activity and an aggressive phenotype, possibly exacerbated by the presence of the I148M PNPLA3 in HepG2 genetic background (3, 7, 9, 11).

Therefore, for the first task of this work, we carried out a detailed morphological and functional characterization of mitochondria in liver biopsies obtained from 28 newly diagnosed MASLD patients (Discovery cohort), recruited at the Metabolic outpatients’ service at Fondazione IRCCS Cà Granda, Policlinico of Milan, attempting to validate the results obtained *in vitro*. Based on prior findings, we paid the attention on mt-shape, number, and size, evaluated by TEM, and on hepatic expression (IHC) of key proteins regulating mt-turnover of MASLD individuals stratified according to *PNPLA3*, *MBOAT7* and *TM6SF2* genetic background (NRV). The hepatic assessment of mt-biogenesis was then validated in a subset of liver biopsies (n=201) and in 7 MASLD-HCC specimens belonging to Validation cohort, totally including 824 biopsy-proven MASLD outpatients, by evaluating *PGC1-α* mRNA levels and mtDNA copies through *D-loop*, marker of mt-mass.

TEM analysis revealed the presence of canonical mt-structures typically giant, swelled, hypodense and with PIs. These features have been observed in both MASLD and other types of chronic hepatitis, indicating that such alterations may appear in a non-specific fashion across liver diseases (19–22). Unexpectedly, a new spectrum of mt-lesions emerged in NRV=3 carriers from the Discovery cohort, who exhibited the most severe MASLD phenotype. We previously described in a separate case report involving a 40-year-old woman with severe hypertriglyceridemia and MASLD, other noncanonical mt-aberrancies, such as *onion-like* mitochondria. Whole-exome sequencing analysis of this patient highlighted a wide range of SNPs implicated in both lipid handling and mt-biological processes, such as *NDUFAF1* and *POLG* (10). Notably, these mt-related genes may not directly alter mt-morphology, supporting that genetic factors involved in lipid metabolism may contribute to mt-morphological alterations, likely through changes in lipid composition (3). These findings were corroborated in our Discovery cohort, in which mt-structural derangements, including double membrane ruptures, irregular cristae arrangements, and mt-vacuolization were exclusively detected in subjects with all three SNPs in *PNPLA3*, *MBOAT7* and *TM6SF2* genes. Co-presence of missense mutations in these genes, also regulating hepatic fat metabolism, may explain the phenotypic variability observed in hepatic mt-structures compared to the other MASLD patients with the similar histological degree.

At present, conflicting results have been reported regarding the expression of mt-markers and mt-content in MASLD. Carabelli and collaborators argued that mitobiogenesis and mtDNA-CN increase in high-fat diet fed rats, while Li et al supported that hepatic mitobiogenesis, boosted through *PGC1-α*, alongside OPA1 overexpression drives tumor growth, invasion, and associated with poor HCC outcomes, thereby sustaining our findings (3, 23–25). Conversely, several studies have reported scarce mt-content and low *PGC1-α* expression in MASLD (21). More consistent are data concerning mitophagy, which suggested that reduced PINK-PRKN signalling is responsible of defective mt-accumulation (26). Here, a significant increase of hepatic mt-number and dimension, quantified by TEM, paralleled by a marked *PGC1-α* nuclear positivity in hepatocytes was found in NRV=3 patients of the Discovery cohort. Mt-mass correlated with the presence of NRV=3 variants at multivariate analysis, adjusted for NAS score, supporting a direct effect of genetics in modulating mt-dynamics. Furthermore, high levels of OPA1 and DRP1, involved in mt-fusion and fission, respectively, were observed in NRV=3 patients, thus accounting for the formation of megamitochondria and the large quantity of mitochondria within liver biopsies. Importantly, although hepatic PINK1 levels were also upregulated in NRV=3 carriers, PRKN was reduced thus implying that the early termination of mitophagy cascade may be at the basis of the the considerable number of ultrastructural abnormalities observed in these patients. Elevated levels of *PGC1-α* transcripts, *D-loop* and mtDNA-CN were even detected in NRV=3 hepatic samples of the Validation cohort and the greatest amount of mtDNA were found in a NRV=3 MASLD-HCC carrier , thereby suggesting that mt-biogenesis is markedly enhanced in individuals with a strong genetic predisposition, leading to an overall increase of mt-biomass.

Although our findings partially diverge from previous reports, there are plausible explanations for these discrepancies. Firstly, the lack of detailed characterization of the entire hepatic mt-lifecycle specifically in MASLD outpatients. Few human evidence has emerged from case studies of subjects undergoing bariatric surgery, who differ from ours for class enrollment criteria and anthropometric parameters that can modulate mt-content (27). Additionally, existing preclinical dietary models don’t fully reflect the human condition or account for genetic variations affecting lipid metabolism. Still, markers of mt-function (i.e. *MT-ND1*) rather than direct mt-mass measurements may potentially lead to inaccurate assessment of mt-content (28).

To deepen the influence of genetics on mt-markers, we correlated TEM and IHC quantifications with the individual *PNPLA3*, *TM6SF2*, and *MBOAT7-TMC4* polymorphisms, an aspect still unknown. I148M PNPLA3 carriers showed higher PGC-1α, OPA1, DRP1, and PINK1 positivity compared to non-carriers, without unvarying PRKN levels and mt- content, supporting a potential compensatory mechanism that maintain total mt-mass unchanged.

The *MBOAT7-TMC4* T allele carriers enhanced PGC-1α and mt-number, but without showing alterations in fusion/fission markers. Intriguingly, the increase in PINK1 +ve areas suggested that the *MBOAT7* variant may affect mt-biomass through the modulation of mitophagy. Instead, *TM6SF2* mutation carriers exhibited the highest mt-number and PGC-1α levels, raised expressions of OPA1, DRP1, and PINK1, and decreased PRKN. Therefore, our results sustained that *MBOAT7* and *TM6SF2* SNPs, in addition to influencing mt-biogenesis, compromise the degradation of these organelles, generating a significant unbalance of hepatic mt-biomass. Importantly, these assumptions were supported by multivariate analysis which revealed that *MBOAT7-TMC4* and *TM6SF2* risk alleles had a greater impact on altering mt-content rather than *PNPLA3* variant, further confirming the previous findings in the genetic *in vitro^-^* models (3, 7, 9, 11).

The second task of this work aimed to translate our observations into clinical practice with the goal to create dynamic risk scores for MASLD monitoring. For this purpose, we move forward towards an extensive literature research on mt-derived biomarkers reported in hepatopathy or metabolic diseases, that could be surrogates of mt-mass and function. Prior studies reported that mtDNA-CN, assessed in PBMCs, increased in diabetic patients and in those with MASLD diagnosis compared to non-MASLD (29–31). Moreover, a huge percentage of mtDNA mutations in *D-loop* region, enhancing mtDNA-CN, negatively associated with HCC clinicopathological profile (32, 33). Similarly, serum ccf-mtDNA fragments derived from respiratory chain raised in MASH patients, particularly in those with significant fibrosis, and were proposed as prognostic tumoral markers (34–36).

Therefore, for mt-content, the choice fell on *D-loop*, regulating mtDNA replication, which was assessed in PBMCs. Instead, a mt-encoded fragment of the OXPHOS (ccf-COXIII) was selected as marker of mt-activity and measured in MASLD serum samples. Both biomarkers were evaluated the Discovery and Validation cohorts and correlated with genetic profile and disease severity (11). We found that NRV=3 carriers of both Discovery/Validation cohorts, showed the highest circulating *D-loop* levels suggesting that as in the liver, this biomarker reflects the increased mt-mass. Moreover, at multivariate analysis, *D-loop* was independently modulated by the presence of *MBOAT7* and *TM6SF2* SNPs rather than by the I148M PNPLA3 variant in the Validation cohort, thereby reinforcing the hypothesis that these genes directly affect mt-dynamism. Moreover, *D-loop* levels positively correlated with MASLD-related metabolic comorbidities (, transaminases and progressively increased with hepatic histological damage in an independent manner, thus suggesting this biomarker may be a relevant pathological sign of MASLD.

Concerning ccf-COXIII, NRV=3 patients augmented the concentration of this marker and, differently from *D-loop*, its levels were associated with all the *PNPLA3*, *MBOAT7*, and *TM6SF2* individual SNPs. Thus, despite the I148M PNPLA3 mutation exerts a minor effect on mt-dynamics compared to *MBOAT7* and *TM6SF2* at-risk genotypes, its role stands out most for mt-dysfunction. Such findings meet confirmation in a recent work proposed by Silva et al, who carried out a metabolome analysis in more than 2000 MASLD participants, revealing positive associations between from TCA metabolites and the I148M PNPLA3 polymorphism (37). Moreover, Caon et al and Luukkonen et al showed that mt- respiratory chain and antioxidant defense decreased in hepatic stellate cells (HSCs) carrying the I148M PNPLA3 variant (38) as well as I148M homozygous patients reduced TCA flux and increased hepatic mt-β-oxidation, redox state and ketogenesis (39).

Notably, NRV=3 patients represent the fraction of subjects with the highest disease severity. Accordingly, although ccf- COXIII was not associated with environmental risk factors and MASLD early stages, its release was extremely high in cirrhotic and HCC subjects, thus representing a potential biomarker for these conditions.

Given the evident link among genetics, mt-biomarkers and disease progression, the final aim of this work focused on developing noninvasive risk scores integrating genetic information, biochemical parameters, metabolic comorbidities, and mt-derived biomarkers in the attempt to overcome existing challenges (12, 13). A significant dilemma is the absence of effective scoring systems for early detection of MASH, a reversible stage in patients. This deficiency may be due to the weakness of novel proposed biomarkers, which may have a pathological meaning but fails alone to predict MASLD progressive forms, to the extent that new diagnostic/prognostic approaches involve their use in combination and/or with other biomarkers of liver damage, such as transaminases (12, 40). PRS have also found their utility for clinical risk stratification in MASLD. However, their predictive power is often limited by low AUC, sensitivity, or specificity values. Additionally, incorporating genetic assessments into existing scores for fibrosis and HCC diagnosis, which combine clinical factors, did not significantly improve the predictive accuracy of the models. (16, 41, 42).

As abovementioned, generative AI tools have made possible to extract clinically relevant information from large dataset, mostly implementing histopathological analysis and further diagnosis of MASLD and HCC through machine and deep learning techniques (43). Similarly, here we provided a totally anonymized dataset to rsGPT-4, which aid to extrapolate defined features (age, BMI, genetic variants, and the two novel mt-biomarkers) and to optimize their combination to get robust prediction models for MASH, fibrosis, and HCC. The rsGPT-4 proposed a RF analysis which allowed to consider the complex interactions and non-linear relationships among the inputted variables. Importantly, for rigorousness and scientific validity, we manually run the RF algorithms in JMP Pro 17 statistical software, collected the relative features importance and gathered them in a linear formula. All the steps were performed to develop the “MAGIC”-named scores, which have in common age, BMI and genetic factors. The MAGIC- scores differ based on the presence of only one mt- biomarker (*D-loop* or ccf-COXIII), both and their relationship with *PNPLA3*, *TM6SF2* and *MBOAT7-TMC4* SNPs. Another major difference is the outcome for which they were generated. In fact, RF analyses were performed by placing MASH, fibrosis, or HCC as final endpoint. By applying the new risk scores to the Validation cohort, the MAGIC-MASH and the MAGIC-Fib including anthropometric, genetic data and the *D-loop* alone resulted in models with the best AUC (73% and 76%, respectively) compared to those with ccf-COXIII alone or both biomarkers, with a discrete sensitivity and specificity. Such findings support that *D-loop* assessment may be sufficient for the early detection of MASH and fibrosis. For HCC, the ccf-COXIII reached an AUC of 81%, suggesting that it is a more effective than *D-loop* (77%). Furthermore, the combination of both biomarkers improved the accuracy up to 86%, without restricting sensitivity and specificity parameters (78.6%-81.5%), and enhancing HCC detection, particularly at a risk score threshold near 30.

To sum up, new hits emerged with this study. As first, mounting evidence is pointing out that genetics cannot be thought of as a marginal part of pathology. Numerous studies have demonstrated how the I148M PNPLA3 mutation accelerates disease progression, affecting both hepatocytes and HSCs metabolism. In keeping with these findings, our data supported the role of this variant in altering hepatic mt-activities but not mt-dynamics, thus adding further insights into how this gene may contribute to liver damage. The rs641738 C>T in *MBOAT7-TMC4* locus and the E167K *TM6SF2*, which are the second major MASLD predictors, are less studied regarding their influence on disease progression. Our findings in MASLD patients, previously supported by the experimental data, revealed that the cumulative presence of 148M PNPLA3, the rs641738 *MBOAT7-TMC4* and the E167K TM6SF2 SNPs increases the risk of advancing from SLD up to HCC. It also highlighted a new aspect related to *MBOAT7* and *TM6SF2* loss-of-functions, which impact on both mt- biogenesis and functions possibly altering hepatic lipid species and organelles’ interplay, a mechanism that may underlie to disease progression.

Secondly, the study dissected the role of two novel biomarkers of mt-origin, whose circulating levels depend on the examined SNPs and MASLD severity. While both markers act as pathological indices, their clinical significance varies: *D-loop*, progressively rising with steatosis, necroinflammation and fibrosis degrees, is highly informative as early MASLD indicator (MASH and fibrosis), while ccf-COXIII is more closely related to cirrhosis and HCC, and more useful for monitoring the worst disease forms. As both mt-biomarkers remarkably increase in HCC cases, their combination in the MAGIC-HCC score provided the best prognostic performance for HCC. Of note, we recognize that developing new risk scores with machine learning methods can be challenging and that AI adoption in clinical settings may face resistance. Nevertheless, AI support allowed this study to pioneer the proposal of risk scores that, for the first time, could non-invasively distinguish between MASH and non-MASH cases. It helped optimize the use of genetic data and circulating biomarkers while considering the causal relationship between them, enabling the generation of scores with an AUC comparable to the existing ones for fibrosis and HCC without significantly compromising sensitivity or specificity. It’s crucial to emphasize that AI is a technological innovation which, if used ethically and with careful consideration of the experts—who remains irreplaceable—can be a turning point, helping clinicians and researchers in refining current methods.

## Supplemental Materials and Methods

### Patients and Methods

#### Discovery Cohort

From 2019 to 2023, n=28 unrelated patients of European descent were consecutively enrolled at the Metabolic Liver Diseases outpatient service at Fondazione IRCCS Cà Granda, Ospedale Maggiore Policlinico Milano (Milan, Italy). Inclusion criteria were the availability of a liver biopsy specimen for suspected MASH or severe obesity, DNA samples, and clinical data. Individuals with excessive alcohol intake (men, >30 g/d; women, >20 g/d), viral and autoimmune hepatitis, or other causes of liver disease were excluded. The study conformed to the Declaration of Helsinki and was approved by the Institutional Review Board of the Fondazione Ca’ Granda IRCCS of Milan and relevant institutions. All participants provided written informed consent. For each patient, the liver biopsy underwent a dedicated inclusion protocol for transmission electron microscopy (TEM) analysis and an expanded panel of immunohistochemistry evaluation of mt-morphology and -dynamics markers. The discovery cohort was stratified according to the number of risk variants (NRV) as previously described (1): 0 for patients who had no risk alleles; 1 for the presence of 1 risk allele heterozygous or homozygous in either *PNPLA3*, *MBOAT7* or *TM6SF2*; 2 for carriers who had 2 risk variants among *PNPLA3*, *MBOAT7* or *TM6SF2* in variable combinations; 3 for subjects carrying all 3 at-risk variants either heterozygous or homozygous. Demographic, anthropometric, and clinical features of the Discovery cohort are shown in **Table S1**.

#### Validation Cohort

The Validation cohort consisted of 824 unrelated MASLD patients of European descent who were mostly recruited at the Metabolic Liver Diseases outpatient service at Fondazione IRCCS Cà Granda, Ospedale Maggiore Policlinico Milano (Milan, Italy). Among the 824 patients, 125 had MASLD-HCC and they were enrolled between January 2008 and January 2015 at the Milan, Udine, and Rome hospitals. The diagnosis of HCC was based on the European Association for the Study of the Liver–European Organization for Research and Treatment of Cancer Clinical Practice Guidelines(2). In absence of a liver biopsy specimen, diagnosis of MASLD-HCC required detection of ultrasonographic steatosis plus at least 1 criterion of the metabolic syndrome. Inclusion and exclusion criteria were the same as described in the Discovery cohort as well as genetic stratification according to the NRV definition (1). All participants provided written informed consent. The demographic, anthropometric, and clinical features of the Validation cohort stratified according to histological clinical phenotype are shown in **Table S2**. Differently from the Discovery cohort, in which mt- morphology/mass and -dynamics markers were assessed by TEM and IHC, we selected n=201 liver biopsies with enough tissues for DNA (n=89), RNA (n=88) and protein extraction (n=24) among MASLD patients belonging to the Validation cohort, to perform gene expression analysis of genes involved in mt-dynamics and for the evaluation of mt-content and mt-damage.

#### Histologic Evaluation

Steatosis was divided into the following 4 categories based on the percentage of affected hepatocytes: 0, 0%–4%; 1, 5%– 32%; 2, 33%–65%; and 3, 66%–100%. Disease activity was assessed according to the nonalcoholic fatty liver disease (NAFLD) activity score (NAS), with systematic evaluation of hepatocellular ballooning and necroinflammation; fibrosis also was staged according to the recommendations of the NAFLD Clinical Research Network^1^. The scoring of liver biopsy specimens was performed by independent pathologists unaware of patient status and genotype. MASH was diagnosed in the presence of steatosis, lobular necroinflammation, and hepatocellular ballooning.

#### Genotyping

The Discovery and Validation cohorts (**Table S1-2**, respectively) were genotyped for the rs738409 C>G (PNPLA3 I148M), rs58542926 C>T (TM6SF2 E167K), and rs641738 C>T MBOAT7 risk variants as previously described^15,16^. Genotyping was performed in duplicate using TaqMan 5’-nuclease assays (QuantStudio 3; Thermo Fisher, Waltham, MA). Results of rs738409, rs58542926, and the rs641738 genetic frequencies were compared with those obtained in non-Finnish European healthy individuals included in the 1000 Genome project^2^

### Transmission Electron Microscopy (TEM)

Mitochondrial morphology, content and ultrastructural defects were assessed in liver biopsies of the Discovery cohort by TEM. Hepatic biopsy was fixed in 2.5% glutaraldehyde in cacodylate buffer pH 7.4, overnight. Afterward, it was postfixed in 2% osmium tetroxide (OsO_4_) for 1h. Finally, the liver specimen was dehydrated with increasing ethanol series, embedded in an Epon resin, and polymerized in an oven at 62°C for 48h. Ultrathin (70–90 nm) sections were collected on nickel grids and observed with a Zeiss EM109^3,4^.

### Immunohistochemistry (IHC)

Attempting to investigate cellular localization, activation and expression of markers of mt-turnover involved in fusion, fission and mitophagy, part of liver biopsy was exploited for immunohistochemistry (IHC). Paraffin-embedded liver biopsies were deparaffined with 100% xylenes and 100% ethanol washes for three times. Antigen retrieval was performed by boiling liver slices in sodium citrate and then permeabilized in 0.3% Triton X-100 for 10 minutes at room temperature (RT). Blocking was executed with 5% Bovine Serum Albumin (BSA) or a mixture of 1X PBS, 0.3% Triton X-100, 10% BSA and 5% Milk according to protein cellular localization (SigmaAldrich, St Louis, MO) for 40-45 minutes at RT. Then, samples were incubated with primary antibodies overnight at 4°C. Anti-rabbit or anti-mouse horseradish-peroxidase– conjugated antibodies were incubated for 2 hours at RT and 3,30-diaminobenzidine (DAB) was provided as chromogen. Nucleus was counterstained with haematoxylin. Finally, samples were mounted with a drop of VectaMount® Express Mounting Medium (Maravai LifeSciences, Inc, San Diego, CA). For each patient, at least n=3 non-overlapping random images were quantified by ImageJ software. Each data point plotted in the bar graph represent the mean value of each measurement *per* patient.

| Primary Antibody<br>(dilution) | Catalogue | Secondary Antibody (dilution) | Catalogue | Blocking solution* |
| --- | --- | --- | --- | --- |
| **Anti-PGC-1α**<br>(1:50) | Cell Signaling<br>#2178 | Anti-IgG Rabbit<br>(1:100) | Cell Signaling<br>#7074 | 1X PBS<br>0.3% Triton X-100<br>10% BSA<br>5% Milk |
| **Anti-mt-Surface**<br>(1:100) | EMD Millipore<br>MAB1273 | Anti-IgG Mouse<br>(1:200) | Cell Signaling<br>#7076 | 1X PBS<br>0.3% Triton X-100<br>10% BSA<br>5% Milk |
| **Anti-MFN1**<br>(1:100) | Abcam<br>ab126575 | Anti-IgG Mouse<br>(1:200) | Cell Signaling<br>#7076 | 5% BSA |
| **Anti-MFN2**<br>(1:100) | Abcam<br>ab56889 | Anti-IgG Mouse<br>(1:200) | Cell Signaling<br>#7076 | 5% BSA |
| **Anti-OPA1**<br>(1:50) | Cell Signaling<br>#67589 | Anti-IgG Rabbit (1:100) | Cell Signaling<br>#7074 | 5% BSA |
| **Anti-DRP1**<br>(1:100) | Cell Signaling<br>#8750 | Anti-IgG Rabbit<br>(1:200) | Cell Signaling<br>#7074 | 1X PBS<br>0.3% Triton X-100<br>10% BSA<br>5% Milk |
| **Anti-PINK1**<br>(1:50) | Cell Signaling<br>#6946 | Anti-IgG Rabbit (1:100) | Cell Signaling<br>#7074 | 5% BSA |
| **Anti-Parkin**<br>(1:100) | Cell Signaling<br>#4211 | Anti-IgG Mouse (1:200) | Cell Signaling<br>#7076 | 5% BSA |

List of primary and secondary antibodies exploited for evaluation of mt-dynamics in n=28 hepatic specimens (Discovery cohort) by IHC. *Reagents for blocking solution were bought from Sigma-Aldrich (St Louis, MO), Cell signaling (Danvers, Massachusetts, USA) and Abcam (Cambridge, UK)

### Gene expression analysis

RNA was extracted from liver biopsies (n=88) using Trizol reagent (Life Technologies-ThermoFisher Scientific, Carlsbad, USA). 1 µg of total RNA was retrotranscribed with a VILO random hexamers synthesis system (Life Technologies- ThermoFisher Scientific, Carlsbad, USA). Quantitative real-time PCR (qRT-PCR) was performed by an ABI 7500 fast thermocycler, using the TaqMan Universal PCR Master Mix (Life Technologies, Carlsbad, USA) or SYBR Green chemistry (Fast SYBR Green Master Mix; Life Technologies, Carlsband, USA). PPARG coactivator 1 alpha (*PGC1-α*, Hs00173304_m1) human taqman probe was bought by Life Technologies-ThermoFisher Scientific (Carlsbad, USA) to assess mt-dynamics. All reactions were delivered in triplicate. Data were normalized to *beta-actin (ACTB)* housekeeping gene and results were expressed as 2^^-ΔCT^ mean value ± standard deviation (SD).

### Western blot

A subset of n=24 MASLD patients enrolled at the Metabolic Liver Diseases outpatient service, Fondazione IRCCS Cà Granda and Ospedale Maggiore Policlinico, Milan, Italy, for whom liver biopsies were available, was stratified according to the number of risk variants (NRV) as follows: 0 for patients who had no risk alleles; 1 for the presence of 1 risk allele heterozygous or homozygous in either PNPLA3, MBOAT7,or TM6SF2; 2 for carriers who had 2 risk variants among PNPLA3, MBOAT7,or TM6SF2 in variable combinations; 3 for subjects carrying all 3 at-risk variants either heterozygous or homozygous. For each group, n=6 hepatic tissues were selected for Western blot analysis. Proteins were extracted from 5 mg of liver samples using RIPA buffer containing 1 mmol/L Na-orthovanadate, 200 mmol/L phenylmethyl sulfonyl fluoride, and 0.02 μg/μL aprotinin. Samples were pooled prior to electrophoretic separation, and all reactions were performed in duplicate. Then, equal amounts of proteins (50 μg) were separated by SDS-PAGE, transferred electrophoretically to nitrocellulose membrane (BioRad, Hercules, CA), and incubated with monoclonal primary antibodies (Total OXPHOS Human WB Antibody Cocktail, Abcam, Cambridge, UK) at 4°C overnight. Total OXPHOS Human WB Antibody Cocktail targets the Complex I subunit NADH:ubiquinone oxidoreductase subunit B8 (NDUFB8), Complex II subunit succinate dehydrogenase complex iron sulfur subunit B (SDHB), Complex III subunit ubiquinol-cytochrome C reductase core protein 2 (UQCRC2), Complex IV subunit mitochondrially encoded cytochrome C oxidase II (COX-II) and ATP synthase F1 subunit alpha (ATP5A), belonging to the oxidative phosphorylation (OXPHOS) system. Finally, samples were incubated with an anti-mouse horseradish peroxidase (HRP)-conjugated antibody for 1 h and Clarity Western ECL substrate (BioRad, Hercules, CA) was used for protein detection. Protein bands from western blot films were quantified through ImageJ software and normalized to SDHB levels, with SDHB used as a mt-housekeeping protein.

#### Quantification of hepatic and circulating mtDNA copy number (CN)

Hepatic mitochondrial DNA (mtDNA) copy number (CN) was assessed in a subgroup of patients (n=89) who belong to the Validation cohort whereas the circulating one was evaluated in PBMCs from both the Discovery (n=28) and Validation (n=824) cohorts. To this purpose, mtDNA was extracted from 5 mg liver biopsies and 200 µl of PBMCs through QIAmp DNA Mini Kit (Manchester, UK). Human samples were resuspended in a Protease K solution, homogenized with a pellet pestel (Sigma-Aldrich, St Louis, MO) and incubated 56°C for 10 minutes to disrupt protein-DNA interactions. Total DNA was trapped onto the QIAamp silica membrane, while contaminants were removed in the flowthrough. Subsequently, DNA was eluted in water and its concentration and quality were assessed by Nanodrop 1000 microvolume 42 spectrophotometer (ThermoFisher Scientific, USA). 5 ng/µl of DNA was used to quantify mtDNA-CN by Quantitative real-time PCR (qRT-PCR, ABI 7500 fast thermocycler, Life Technologies, Carlsbad, USA). Specifically, we measured *D-loop* region, the replication start site of the mtDNA, through TaqMan Copy Number Assay (MT-7S, ThermoFisher #Hs02596861_s1). RNAse-P (ThermoFisher #4401631), a sequence known to exist in two copies in human genome, was used as a reference gene. Median ΔCT per assay value was used as calibrator. mtDNA copies were predicted by analyzing plates through CopyCaller Software 2.0 (ThermoFisher Scientific, USA). *D-loop* expression was calculated with 2^^-ΔΔCT^ and its skewed distribution was logarithmically transformed for the statistical analysis.

#### Measurement of cell-free circulating (ccf) mtDNA

Ccf-mtDNA release was assessed in both Discovery (n=28) and Validation (n=824) cohorts. ccf-mtDNA was isolated from 200 µl of serum samples through QIAmp DNA Mini Kit (Manchester, UK) and extracted as described above. Ccf-mtDNA concentration quality was measured by Nanodrop 1000 microvolume 42 spectrophotometer (ThermoFisher Scientific, USA). 20 ng of ccf-mtDNA and PowerUp SYBR Green Master Mix were exploited for the RT-qPCR assay with ABI 7500 fast thermocycler. The following primers were designed for amplifying the mitochondrially-encoded cytochrome C oxidase III (ccf-COXIII) fragment, a coding subunit of the mt-respiratory chain: forward 5’- TGACCCACCAATCACATGC -3’ and reverse 5’- ATCACATGGCTAGGCCGGAG -3’. Ccf-COXIII serum concentration were obtained by interpolating data to a DNA standard curve with the following points: 73.000 picograms (pg), 36.000 pg, 18.250 pg, 10.000 pg, 5.000 pg and 1.000 pg. Data were expressed as pg/unit and its skewed distribution was logarithmically transformed for the statistical analysis.

#### OpenAI’s GPT-4 support for mtDNA sequencing analysis and risk score development

GPT-4 is an advanced language model by OpenAI, skilled in generating human-like text and understanding complex queries. In coding, it facilitates code generation and debugging, while in statistical analysis, it assists in data interpretation and model selection (https://doi.org/10.48550/arXiv.2303.08774). In this study, the OpenAI GPT-4 was exploited to facilitate the construction of a novel risk score for MASLD diagnosis and/or progression. Stringent anonymization measures were applied to the dataset prior to analysis, removing all direct personal identifiers to safeguard patient confidentiality and adhere to privacy regulations. Regarding mtDNA analysis, we used a customized GPT-4 environment through GPT-builder (https://help.openai.com/en/articles/8770868-gpt-builder), specifically enhanced for advanced coding, statistical analysis and processing large dataset (labelled as risk score GPT-4, rsGPT-4). rsGPT-4 identified key columns corresponding to age, BMI, NRV, *D-loop*, and ccf-COXIII, as essential parameters of interest. Informed by the project’s context, which indicated genetic modulation of mt-biomarker levels, rsGPT-4 suggested the integration of interaction terms between these biomarkers and the genetic risk variables (NRV). Results were meticulously and manually validated in JMP Pro 17 (SAS, Cary, NC).

Endorsing the application of a machine learning approach, rsGPT-4 recommended a Random Forest (RF) model for its proficiency in handling complex datasets and determining predictor importance. Conducted via JMP Pro 17, the RF model—comprising 1000 trees with 3 features sampled at each node—yielded relative feature importance and identified significant interactions between the variables included in the analysis (age, BMI, NRV, *D-loop*, and ccf-COXIII). Through this approach, we developed 3 separated risk scores with the aim to identify the presence of MASH, fibrosis and/or HCC, respectively, named as <u>M</u>itochondrial, <u>A</u>nthropometric, and <u>G</u>enetic Integration with <u>C</u>omputational intelligence (**MAGIC**) for MASH (**MAGIC-MASH**), fibrosis (**MAGIC-Fib**) and HCC (**MAGIC-H**). Furthermore, the contribution of mt-derived biomarkers was assessed by constructing ad hoc scores which included either D-loop alone, ccf-COXIII alone or both. Details on relative features importance and gathered formulas are reported below:

**MAGIC-MASH score**

- ***D-loop*** = (Age*0.284) + (BMI*0.3361) + (NRV=3*0.0758) + (logD-loop*0.1546) + (Interaction_logD- loop/NRV=3*0.1495)
- **Ccf-COXIII** = (Age*0.3566) + (BMI*0.2989) + (NRV=3*0.0736) + (logCOXIII*0.1401) + (Interaction_ logCOXIII/NRV=3*0.1308)
- **Both** = (Age*0.2835) + (BMI*0.1934) + (NRV=3*0.0593) + (logD-loop*0.1085) + (logCOXIII*0.1342) + (Interaction_logD-loop/NRV=3*0.1085) + (Interaction_ logCOXIII/NRV=3*0.105)

**MAGIC-Fib score**

- ***D-loop*** = (Age*0.3247) + (BMI*0.2767) + (NRV=3*0.0787) + (logD-loop*0.1772) + (Interaction_logD- loop/NRV=3*0.1428)
- **Ccf-COXIII** = (Age*0.3458) + (BMI*0.2996) + (NRV=3*0.065) + (logCOXIII*0.1332) + (Interaction_ logCOXIII/NRV=3*0.1563)
- **Both** = (Age*0.2785) + (BMI*0.2229) + (NRV=3*0.0494) + (logD-loop*0.1235) + (logCOXIII*0.1003) + (Interaction_logD-loop/NRV=3*0.111) + (Interaction_ logCOXIII/NRV=3*0.1145)

**MAGIC-H score**

- ***D-loop*** = (Age*0.3427) + (BMI*0.2589) + (NRV=3*0.08) + (logD-loop*0.1678) + (Interaction_logD- loop/NRV=3*0.1506)
- **Ccf-COXIII** = (Age*0.3529) + (BMI*0.2692) + (NRV=3*0.0758) + (logCOXIII*0.1753) + (Interaction_logCOXIII/NRV=3*0.1265)
- **Both** = (Age*0.2835) + (BMI*0.1934) + (NRV=3*0.0593) + (logD-loop*0.1085) + (logCOXIII*0.1342) + (Interaction_logD-loop/NRV=3*0.1085) + (Interaction_ logCOXIII/NRV=3*0.105).

Performance of the risk scores was rigorously evaluated using the AUC-ROC curve, with additional assessments of accuracy, sensitivity, specificity and cut-offs conducted via MedCalc software (MedCalc Software Ltd, Belgium). MAGIC-MASH, MAGIC-Fib and MAGIC-H subcategories with quartiles were applied in our dataset and nominal logistic regression analysis was run to assess the odds ratio.

#### Statistical analysis

For descriptive statistics, continuous variables were reported as means and SD or as the median and interquartile range for highly skewed biological variables. Variables with skewed distribution were logarithmically transformed before analyses. Differences between groups were calculated by one-way nonparametric ANOVA (Kruskal–Wallis), followed by post hoc t-test (two-tailed) when two groups were compared, or the Dunn’s multiple comparison test when multiple groups were compared, adjusted for the number of comparisons. P values < 0.05 were considered statistically significant. Statistical analyses were performed using JMP Pro 17 (SAS, Cary, NC) and Prism software (version 6, GraphPad Software, San Diego, CA).

## Supplemental Results

### MASLD severity modulates mitochondrial morphology and content in the Discovery cohort

We firstly assessed the impact of disease severity on mt-morphology, evaluated by TEM, and mt-dynamics by IHC. The Discovery cohort (n=28, **Table S1**) was stratified according to NAS score. Liver histology showed that 16/28 of cases (57.1%) had a severe disease (NAS≥5), while 4/28 (14.3%) and 8/28 (28.6%) presented a mild (NAS=1-2) or a moderate (NAS=3-4) disease activity, respectively.

As concerns mitochondria, MASLD patients with NAS≥5 displayed on TEM a higher mt-content compared to those with mild/moderate disease (**Figure S1A-B,** p=0.0009 at ANOVA, adj p=0.02 *vs* mild and adj p=0.003 *vs* moderate). At multivariate analysis adjusted for confounding factors potentially modulating mt-mass as age and BMI, the hepatic mt- content associated with a severe NAS (**Table 1**), supporting that disease progression significantly impact on mt- biogenesis.

Moreover, enlarged matrix granules were observed in MASLD patients with moderate NAS, while paracrystalline inclusions (PIs) and formation of giant mitochondria (GM) with PIs represented the major mt-morphological defects observed in MASLD subjects with NAS≥5 (**Figure S1A**). Consistently, measurement of mt-perimeter and area highlighted that severe MASLD subjects presented a larger mt-size compared to those quantified in patients with mild/moderate NAS (**Figure S1C-D**, p<0.0001 at ANOVA; perimeter: adj p=0.002 *vs* mild and adj p=0.0006 *vs* moderate; area: p=0.003 *vs* mild and adj p=0.00002 *vs* moderate), thereby supporting that, despite several signs of mt-damage may arise since early MASLD stages, the greatest effects on mt-structures emerged in subjects with progressive MASLD.

---

**Table S1.**
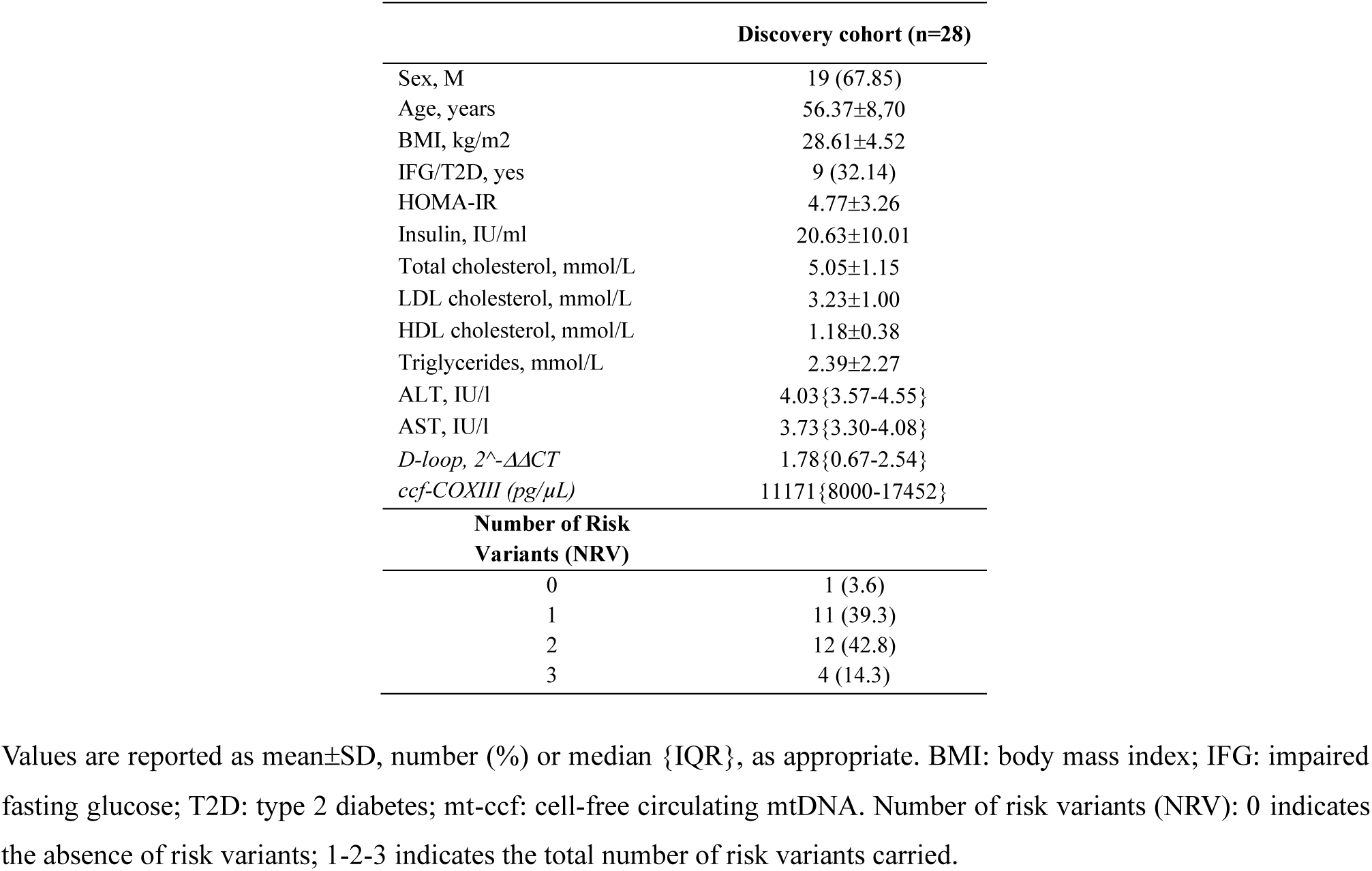
Demographic, anthropometric, and clinical features of n=28 biopsied MASLD outpatients (Discovery cohort)

**Table S2.**
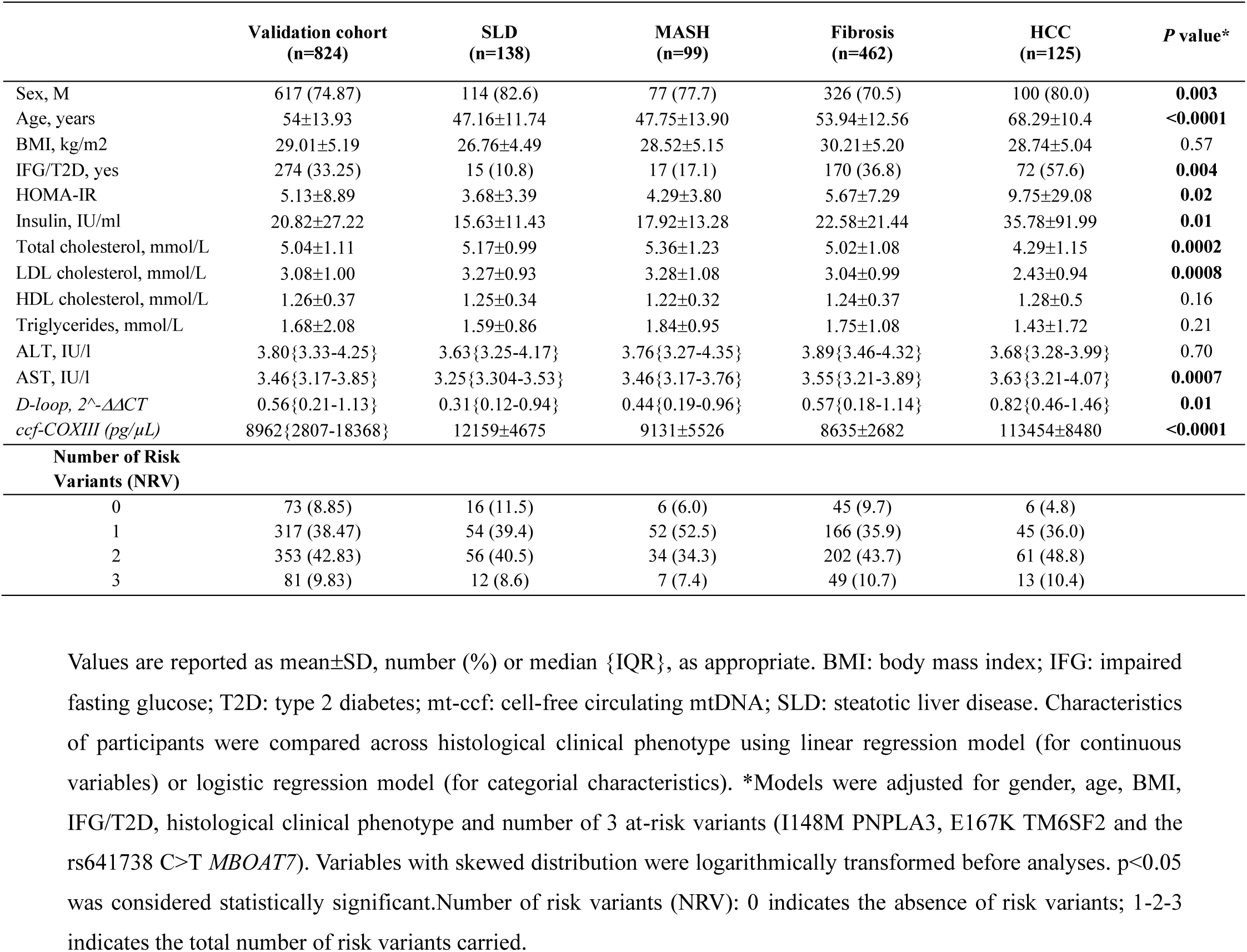
Demographic, anthropometric, and clinical features of n=824 biopsied MASLD outpatients (Validation cohort) stratified according to histological clinical phenotype.

**Table S3.** Generalized linear model and nominal logistic regression analysis correlating circulating *D-loop* levels with anthropometric and biochemical data in the Validation cohort (n=824)

| BMI (Kg/m<sup>2</sup>) |  |  | IFG/T2D, yes |  |
| --- | --- | --- | --- | --- |
| | $\beta$ {95% CI} | *P* value | OR {95% CI} | *P* value |
| Sex, M | -2.97{-3.40--2.45} | <0.0001 | 1.64{1.10-2.45} | 0.01 |
| Age, years | -0.18{-0.21--0.14} | <0.0001 | 1.07{1.05-1.09} | <0.0001 |
| BMI, kg/m<sup>2</sup> | / | / | 1.11{1.07-1.15} | <0.0001 |
| IFG/T2D, yes | 1.42{0.89-1.95} | 0.01 | / | / |
| NRV=3 | -0.71{-1.27-0.15} | <0.0001 | 1.11{0.89-1.38} | 0.34 |
| *D-loop* (log) | 0.49{0.20-0.78} | **0.0008** | 1.22{1.05-1.42} | **0.006** |
| HOMA-IR |  |  | AST (IU/l) |  |
| | $\beta$ {95% CI} | *P* value | $\beta$ {95% CI} | *P* value |
| Sex, M | 0.04{-0.03-0.10} | 0.26 | -0.06{-0.11--0.02} | 0.002 |
| Age, years | 0.003 {-0.001-0.008} | 0.13 | -0.0008{-0.0004-0.002} | 0.61 |
| BMI, kg/m<sup>2</sup> | 0.05{0.04-0.06} | <0.0001 | -0.01{-0.02--0.005} | 0.0005 |
| IFG/T2D, yes | 0.23{0.15-0.30} | <0.0001 | 0.08{0.03-0.12} | 0.0007 |
| NRV=3 | 0.30{-0.04-0.10} | 0.35 | 0.006{-0.04-0.05} | 0.80 |
| *D-loop* (log) | 0.01{-0.03-0.05} | 0.59 | 0.04{0.008-0.07} | **0.01** |
| Lactate (mmol/L) |  |  | LDH (mmol/L) |  |
| | $\beta$ {95% CI} | *P* value | $\beta$ {95% CI} | *P* value |
| Sex, M | -0.03{0.13-0.05} | 0.39 | -0.11{-0.22-0.0008} | 0.05 |
| Age, years | 0.002{-0.004-0.008} | 0.47 | 0.01{0.004-0.02} | 0.002 |
| BMI, kg/m<sup>2</sup> | -0.01{-0.028-0.004} | 0.13 | 0.006{-0.01-0.02} | 0.50 |
| IFG/T2D, yes | 0.14{0.05-0.23} | 0.001 | -0.0006{-0.11-0.10} | 0.90 |
| NRV=3 | -0.02{-0.11-0.07} | 0.62 | -0.01{-0.12-0.09} | 0.80 |
| *D-loop* (log) | 0.07{0.01-0.14} | **0.017** | 0.12{0.03-0.20} | **0.004** |

Values are reported as mean±SD, number (%) or median {IQR}, as appropriate. BMI: body mass index; IFG/T2D: impaired fasted glucose/ type 2 diabetes; NAS: MASLD activity score; NRV: number of risk variants. Ordinal logistic regression analysis was adjusted for sex, age, BMI, IFG/T2D, cumulative presence of rs738409 C>G in *PNPLA3* (I148M), the rs641738 C>T in *MBOAT7-TMC4* locus and the rs58542926 C>T in *TM6SF2* (E167K) and circulating *D-loop*, which was logarithmically transformed. Variables with skewed distribution were logarithmically transformed before analyses (HOMA-IR, AST, lactate and LDH). *p*<0.05 was considered statistically significant.

**Table S4.**
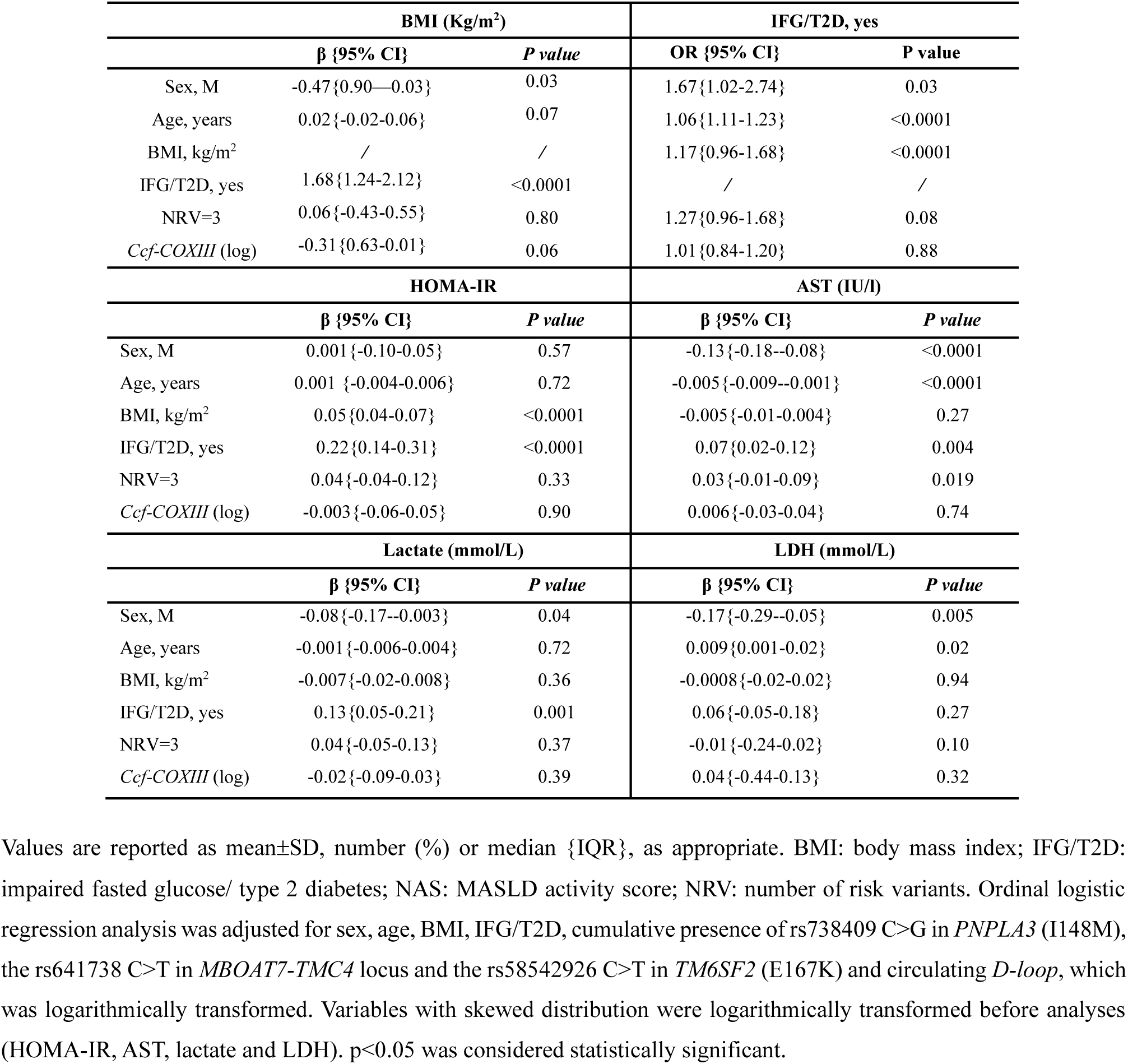
Generalized linear model and nominal logistic regression analysis correlating serum ccf-COXIII levels with anthropometric and biochemical data in the Validation cohort (n=824)

**Table S5.**
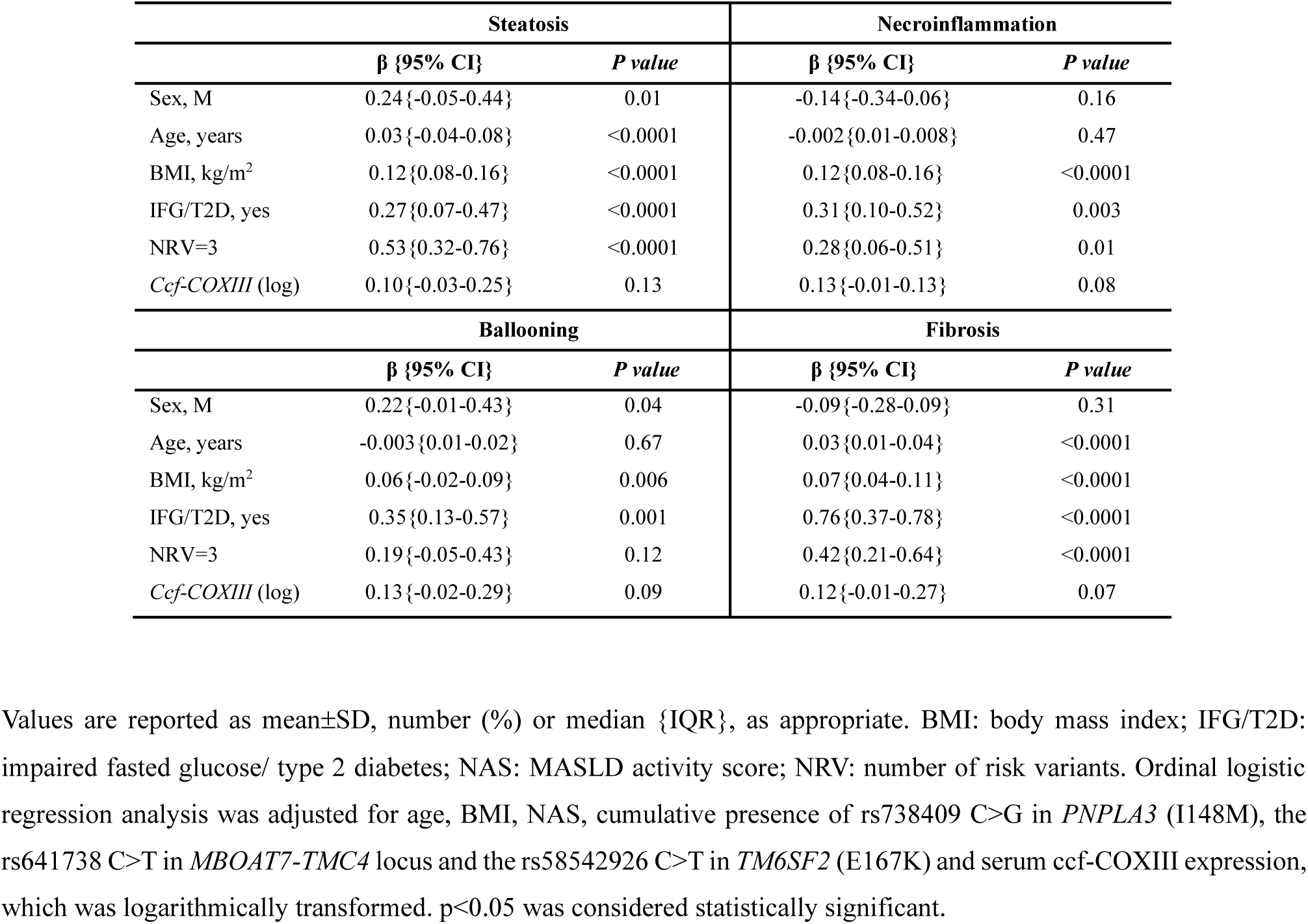
Ordinal logistic regression analysis correlating serum ccf-COXIII levels with histological damage in the Validation cohort (n=824)

| Steatosis |  |  | Necroinflammation |  |
| --- | --- | --- | --- | --- |
| | $\beta$ {95% CI} | *P* value | $\beta$ {95% CI} | *P* value |
| Sex, M | 0.24{-0.05-0.44} | 0.01 | -0.14{-0.34-0.06} | 0.16 |
| Age, years | 0.03{-0.04-0.08} | <0.0001 | -0.002{0.01-0.008} | 0.47 |
| BMI, kg/m<sup>2</sup> | 0.12{0.08-0.16} | <0.0001 | 0.12{0.08-0.16} | <0.0001 |
| IFG/T2D, yes | 0.27{0.07-0.47} | <0.0001 | 0.31{0.10-0.52} | 0.003 |
| NRV=3 | 0.53{0.32-0.76} | <0.0001 | 0.28{0.06-0.51} | 0.01 |
| *Ccf-COXIII* (log) | 0.10{-0.03-0.25} | 0.13 | 0.13{-0.01-0.13} | 0.08 |
| Ballooning |  |  | Fibrosis |  |
| | $\beta$ {95% CI} | *P* value | $\beta$ {95% CI} | *P* value |
| Sex, M | 0.22{-0.01-0.43} | 0.04 | -0.09{-0.28-0.09} | 0.31 |
| Age, years | -0.003{0.01-0.02} | 0.67 | 0.03{0.01-0.04} | <0.0001 |
| BMI, kg/m<sup>2</sup> | 0.06{-0.02-0.09} | 0.006 | 0.07{0.04-0.11} | <0.0001 |
| IFG/T2D, yes | 0.35{0.13-0.57} | 0.001 | 0.76{0.37-0.78} | <0.0001 |
| NRV=3 | 0.19{-0.05-0.43} | 0.12 | 0.42{0.21-0.64} | <0.0001 |
| *Ccf-COXIII* (log) | 0.13{-0.02-0.29} | 0.09 | 0.12{-0.01-0.27} | 0.07 |

Values are reported as mean ± SD, number (%) or median {IQR}, as appropriate. BMI: body mass index; IFG/T2D: impaired fasted glucose/ type 2 diabetes; NAS: MASLD activity score; NRV: number of risk variants. Ordinal logistic regression analysis was adjusted for age, BMI, NAS, cumulative presence of rs738409 C>G in *PNPLA3* (I148M), the rs641738 C>T in *MBOAT7-TMC4* locus and the rs58542926 C>T in *TM6SF2* (E167K) and serum ccf-COXIII expression, which was logarithmically transformed. $p<0.05$ was considered statistically significant.

**Figure S1:**
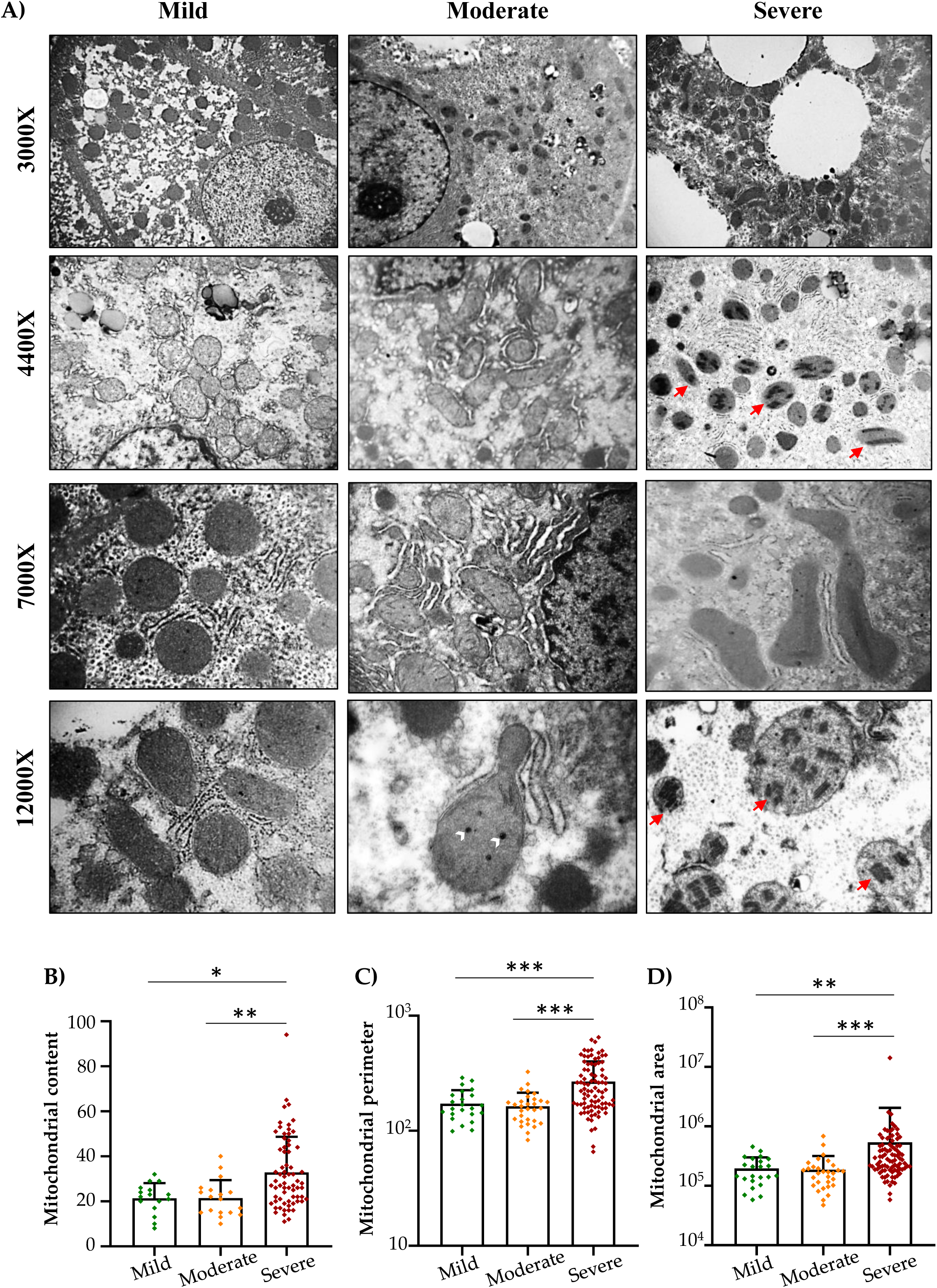
MASLD severity modulates mt-content and morphology in the Discovery cohort. **A**) Representative TEM images showing mt-mass (on top, 3000X magnification) and mt-damage (on bottom; 4400X and 12000X magnification) in NAFLD patients with a mild, moderate or severe disease. White arrowheads showed enlarged matrix granules, while red arrows mark paracrystalline inclusions (PIs). **B**) mt-content was calculated by counting mitochondria in a range of 4- 6 random images per patient (3000X magnification) by ImageJ. **C-D**) Mt-perimeter and area were measured through ImageJ software by tracing the boundary of mitochondria at 7000X magnification. Each data point in the bar graphs represents the mt-number (**B**) or mt-size (**C-D**) measured by ImageJ software in a range of 4-6 random images per patient.

**Figure S2:**
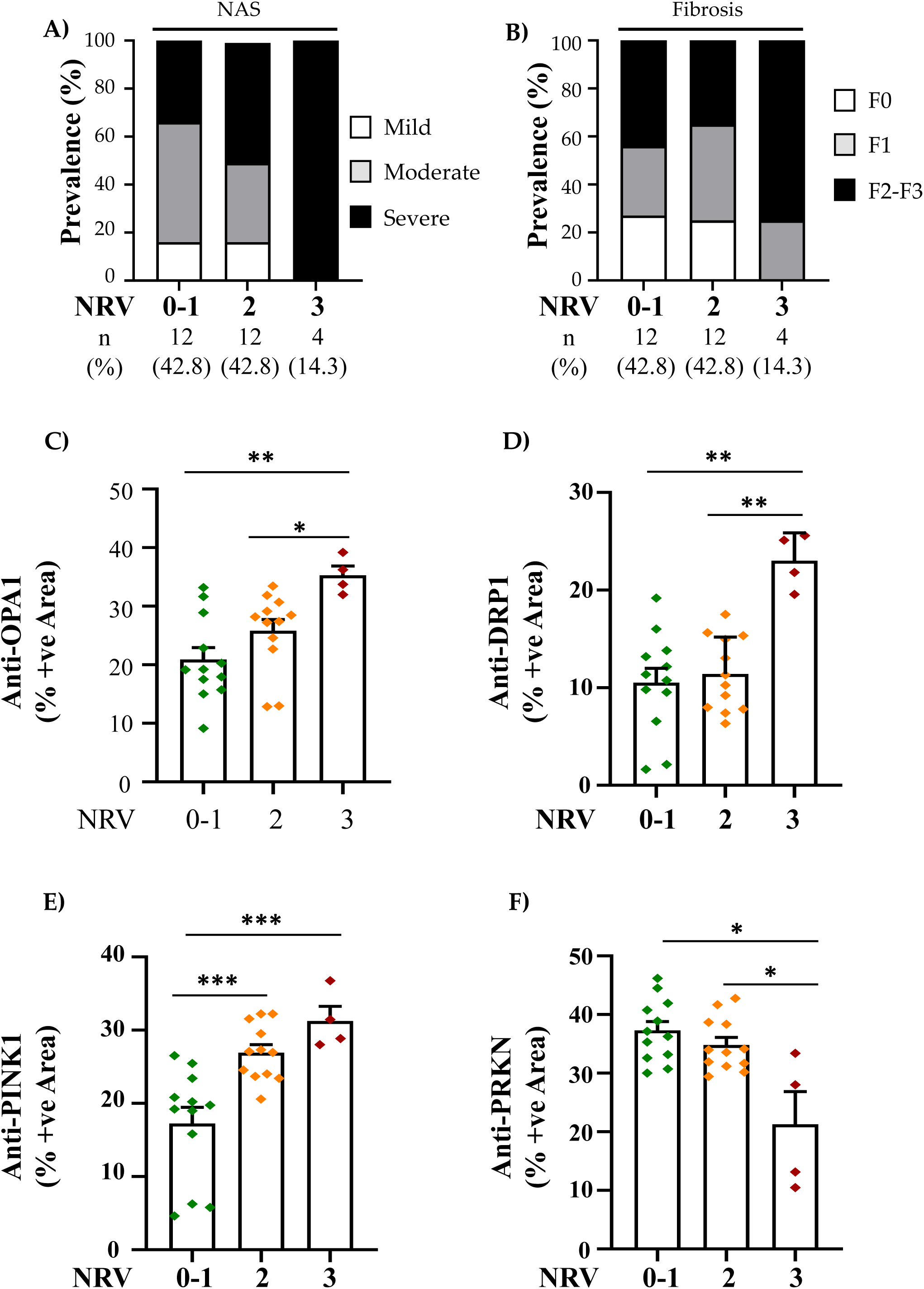
Histological features of MASLD patients of the Discovery cohort and IHC quantification. **A-B)** Contingency analysis showing the distribution (%) of MASLD patients across NRV stratification, falling into NAS subcategories (mild, moderate, severe) and fibrosis stages (F0, F1 and F2-4). **C-F**) Measurement of the area fraction per image, representing the positivity of mt-markers (OPA1, DRP1, PINK1, PRKN; brown-colored cytoplasm), was performed by splitting the RGB channels in red, green and blue separate components, setting a threshold and quantifying the brown intensity through ImageJ software. Each data point in the bar graphs represents the mean value of measured IHC images (a range of 4-6 random non-overlapping photos per patient, 40X magnification).

**Figure S3:**
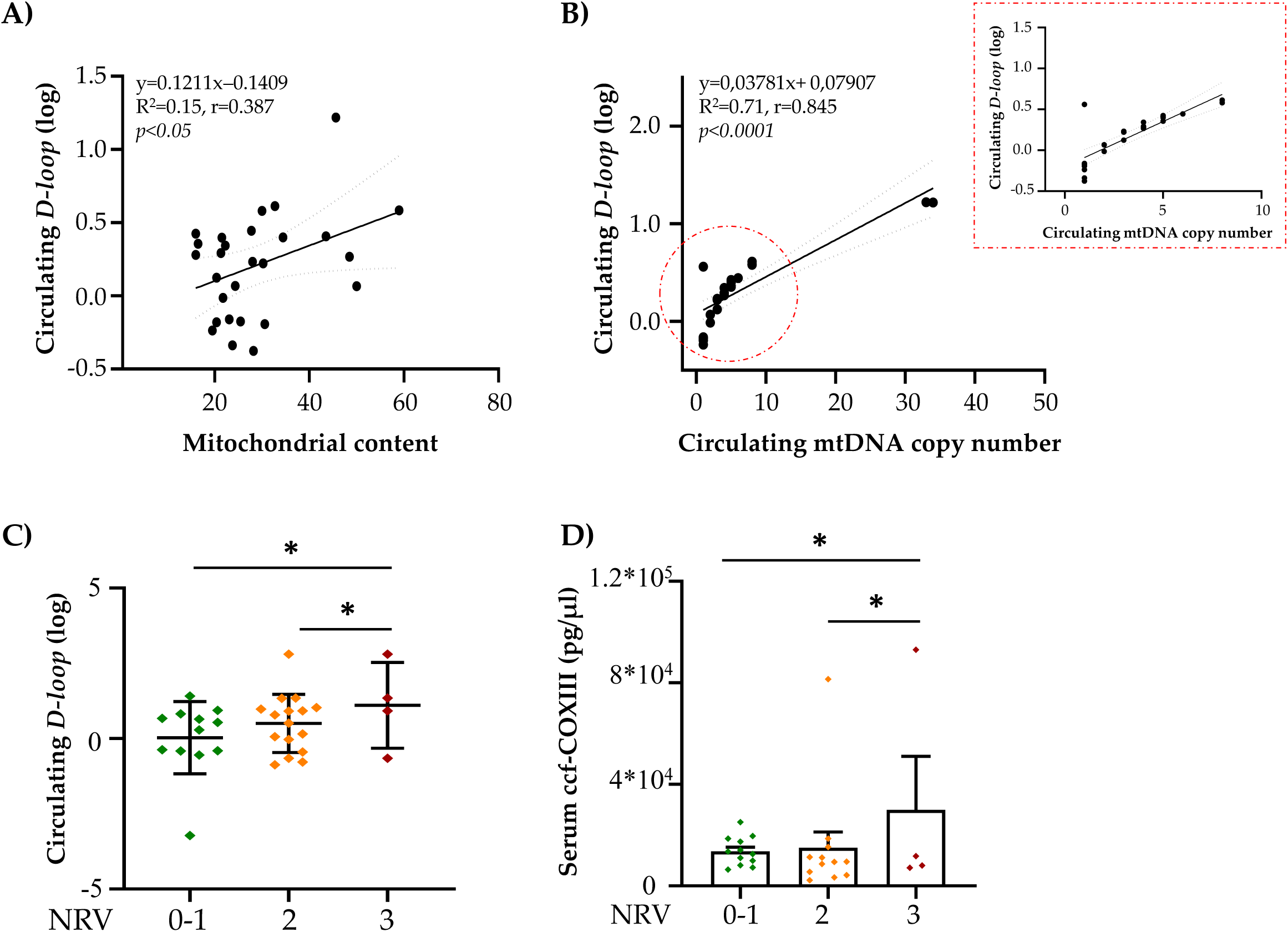
Evaluation of mt-derived biomarkers in MASLD patients of the Discovery cohort. **A**-**B**) Bivariate analysis correlating circulating *D-loop* levels with hepatic mt-content measured by TEM, and with predicted mtDNA copy number (CN) into the bloodstream. **C-D**) Fluctuation of circulating *D-loop* and ccf-COXIII, measured in PBMCs and serum samples respectively, in MASLD patients stratified according to NRV.

**Figure S4:**
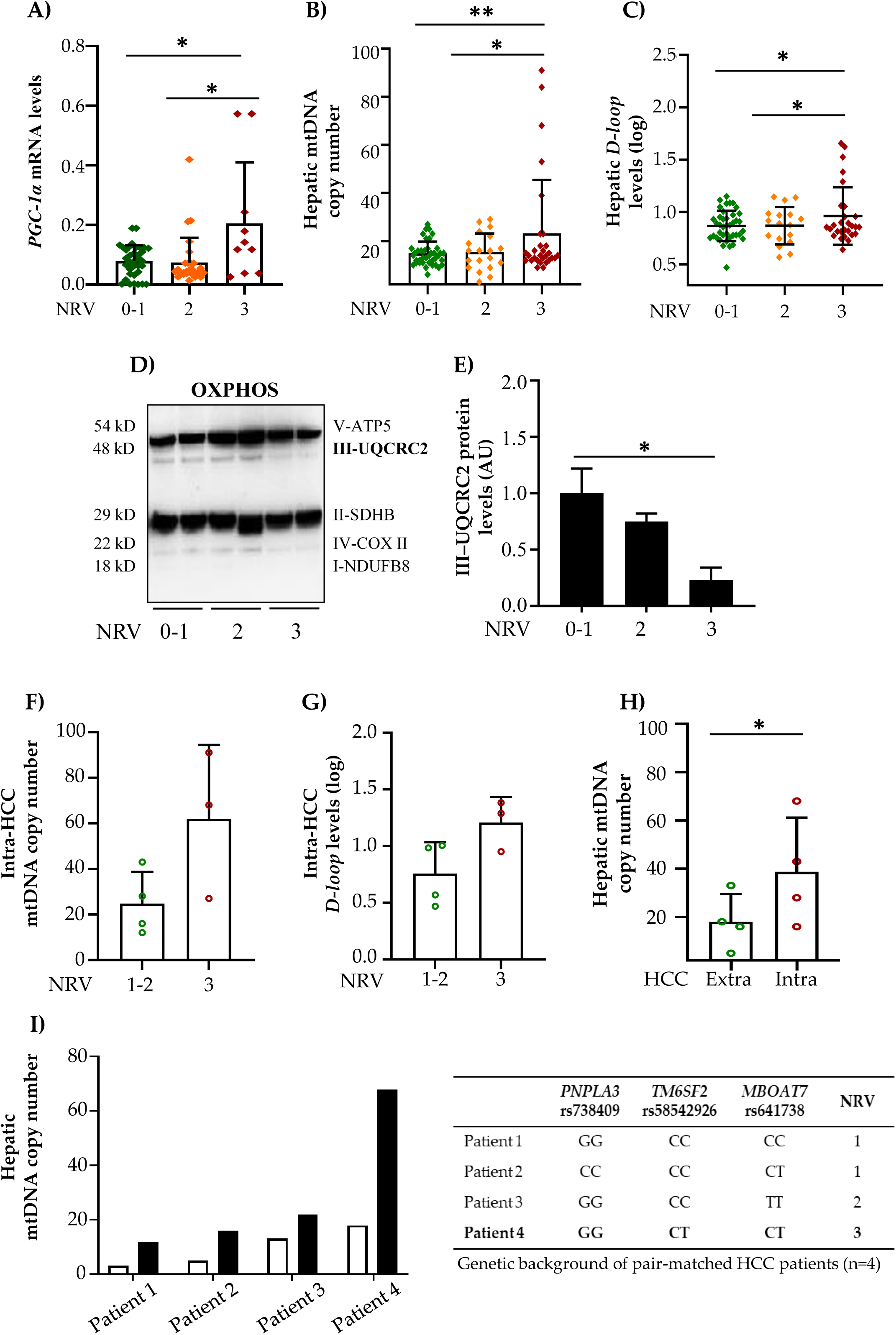
Hepatic assessment of mt-dynamics in subsets of MASLD patients belonging to the Validation cohort and in HCC resections. **A**) *PGC-1α* mRNA levels assessed in 88 liver biopsies of MASLD patients stratified according to NRV. Data were normalized to *beta-actin* (*ACTB*) housekeeping gene and results were expressed as 2^^-ΔCT^ mean value ± standard deviation (SD). **B**-**C**) mtDNA-CN and *D-loop* levels were measured in 89 liver biopsies of MASLD patients stratified according to NRV. mtDNA-CN were predicted through CopyCaller Software 2.0. *D-loop* levels were normalized on *RNAse-P* reference gene. Data are shown as 2^^-ΔΔCT^ mean value ± SD. **D-E**) Western blot analysis of OXPHOS complexes performed in 24 liver biopsies of MASLD patients, pooled and stratified according to NRV. Complex III (UQCRC2) protein levels were quantified through ImageJ software and normalized on Complex I (NADH) mitochondrial housekeeping protein. Data are shown as arbitrary unit (AU-fold increase) ± SD. **D-E**) Measurement of mtDNA-CN and *D-loop* in 7 MASLD-HCC resections. **H**) Comparison of mtDNA-CN between intra- and extra-tumoral tissues. **I**) Comparison of intra- and extra-tumoral mtDNA-CN detected in 4 MASLD-HCC patients stratified according to NRV.

**Figure S5:**
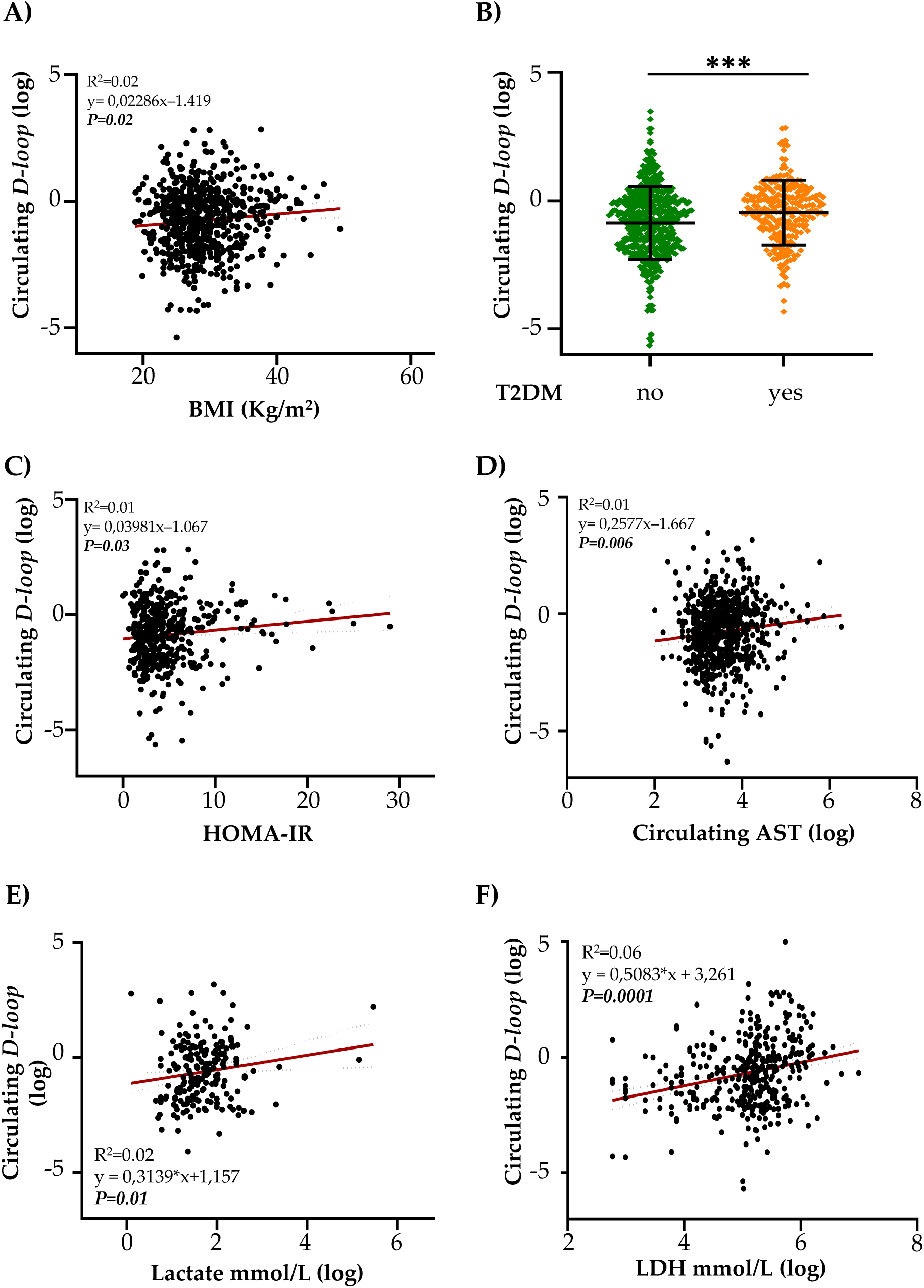
Circulating assessment of D-loop and its associations with metabolic and biochemical traits in the Validation cohort. Bivariate analysis showing a positive correlation among circulating *D-loop* levels and body mass index (BMI) (**A**), type 2 diabetes (T2DM) (**B**), HOMA-IR (**C**), aspartate aminotransferase (AST) (**D**), lactate (**E**) and lactate dehydrogenase (LDH) (**F**).

